# NeighbourNet: Scalable cell-specific co-expression networks for granular regulatory pattern discovery

**DOI:** 10.1101/2025.03.27.645629

**Authors:** Yidi Deng, Jiadong Mao, Jarny Choi, Kim-Anh Lê Cao

## Abstract

Gene regulatory networks (GRNs) provide a fundamental framework for understanding the molecular mechanisms that govern gene expression. Advances in single-cell RNA sequencing (scRNA-seq) have enabled GRN inference at cellular resolution; however, most existing approaches rely on predefined clusters or cell states, implicitly assuming static regulatory programs and potentially missing subtle, dynamic variation in regulation across individual cells. To address these limitations, we introduce NeighbourNet (NNet), a method that constructs cell-specific co-expression networks. NNet first applies principal component analysis to embed gene expression into a low-dimensional space, followed by local regression within each cell’s k-nearest neighbourhood (KNN) to quantify co-expression. This approach improves computational efficiency and stabilises co-expression estimates, mitigating challenges posed by small sample sizes in KNN regression and the inherent noise and sparsity of scRNA-seq data. Beyond co-expression, NNet supports scalable downstream analyses, including (i) clustering and aggregating cell-specific networks into meta-networks that capture primary co-expression patterns, and (ii) integrating prior knowledge to annotate co-expression and infer active signalling interactions at the individual cell level. All functional modules of NNet are implemented with an efficient algorithm that enables application to large-scale single-cell datasets. We demonstrate NNet’s effectiveness through three case studies on transcription factor activity prediction, early haematopoiesis, and tumour microenvironments. Provided as an R package, NNet offers a novel framework for exploring cellular variation in co-expression and integrates seamlessly with existing single-cell analysis workflows.

## 1 Introduction

Gene regulatory networks (GRNs) are essential frameworks for understanding the complex molecular interactions that govern gene expression in biological systems. Central to gene regulation is the binding of transcription factors (TFs) to specific DNA sequences, which can either activate or repress target gene expression. This regulation directs crucial cellular processes such as differentiation, proliferation, and responses to environmental stimuli. The emergence of next-generation sequencing, particularly in transcriptomics in the early 2000s, has enabled high-throughput quantification of gene expression. Since then, substantial research efforts have been dedicated to developing statistical GRN inference methods that leverage associations in gene expression (co-expression) to decipher gene regulation [Huynh-Thu et al., 2010, Langfelder and Horvath, 2008]. These advances offer deep insights into cellular function and have contributed to the identification of key regulators of disease development, providing potential avenues for therapeutic intervention.

Single-cell RNA sequencing (scRNA-seq) has provided an unprecedented view of gene regulation by capturing gene expression profiles at the level of individual cells [Badia-i Mompel et al., 2023]. Cells are often grouped into clusters based on their expression profiles to represent predefined cell states, and GRNs are inferred by measuring co-expression within these clusters to recover cell state-specific regulatory programs [Chan et al., 2017, Zhang et al., 2023, Morabito et al., 2023, Kamimoto et al., 2023]. While clustering-based GRN inference methods have been the most widely investigated, they have a major limitation in assuming that cells within a cluster share similar and discrete regulatory programs. This assumption can mask subtle regulatory changes within each cluster and overlook fine-grained regulatory interactions that occur during cell state transitions. In addition, the accuracy of clustering-based methods relies heavily on the accuracy of cell clustering, meaning that poorly defined clusters can lead to biased or oversimplified findings.

To overcome these limitations, researchers have developed cell-specific GRN inference methods that measure co-expression within the local neighbourhood of each cell. These methods can be broadly classified into two categories based on how neighbouring cells are defined. Some methods define neighbouring cells based on their similarity in expression profiles [Zhang et al., 2022, Dai et al., 2019, Wang et al., 2021]. Co-expression measures in these methods are typically based on correlation measures, meaning that they do not necessarily indicate gene regulation. Other methods define neighbouring cells along a developmental timeline, which can be inferred through pseudotime analysis [Zhang and Stumpf, 2023, Wang et al., 2023]. In this approach, co-expression measures are time-dependent, potentially capturing causal (regulatory) relationships by associating gene expression at successive points in the trajectory. Despite their advantages, time-dependent methods encounter scalability issues. Inferring networks for thousands of cells is computationally intensive and is often restricted to studying only TF interactions, limiting the biological depth of the network. Moreover, for both types of cell-specific methods, the intrinsic sparsity and noise in scRNA-seq data can undermine the stability of co-expression estimates within small neighbourhoods, posing further challenges for reliable network inference [Pratapa et al., 2020, Nguyen et al., 2021].

To address these challenges, we present NeighbourNet (NNet), a novel method that uses k-nearest neighbours (KNN) principal component (PC) regression on gene expression data to efficiently construct robust, cell-specific co-expression networks (Figure 1). Rather than directly interpreting these networks as regulatory, NNet incorporates a scalable downstream analysis framework consisting of: meta-network analysis to uncover common co-expression patterns, meta-TF analysis to identify gene modules, and prior knowledge integration to facilitate cell-specific inference of active gene regulation and upstream signalling pathways (USPs). Our KNN-PC regression approach offers several advantages. First, PCA reduces the high dimensionality of gene expression and boosts the computational efficiency of regression-based co-expression inference. Second, embedding genes into PC space effectively imputes the sparse gene expression, enhancing the stability and accuracy of co-expression measures. Therefore, the NNet framework enables the construction of large-scale cell-specific networks that are both efficient to compute and robust against noise.

**Figure 1.**
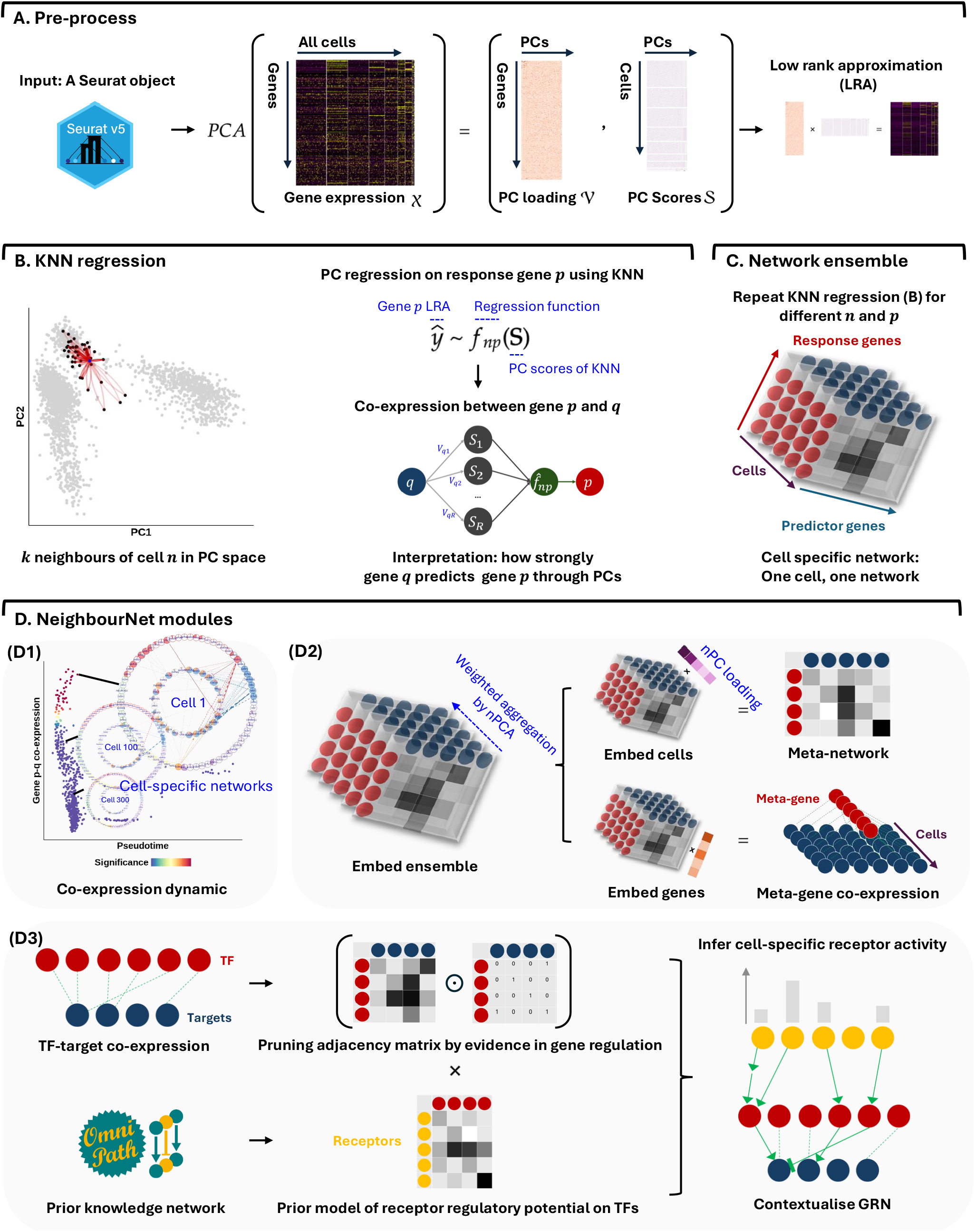
NeighbourNet (NNet) workflow for inferring cell-specific co-expression networks and integrating prior knowledge.. **A. Pre-processing**: Single-cell gene expression data are subject to principal component analysis (PCA) to capture the major variation. A low-rank approximation (LRA) then reconstructs the expression matrix using the PCs. A weighted k-nearest neighbour (KNN) graph that defines each cell’s local neighbourhood in the PC space is constructed. **B. Neighbourhood Regression**: For each cell, NNet performs regression within the cell’s KNN using the LRA-derived gene expression as the response and PCs as predictors. This process quantifies co-expression between genes by measuring how ‘predictor genes’ (those used to embed PCs) contribute to predict each response in the PC space. Repeating this for multiple response genes yields cell-specific co-expression networks. **C. Network Ensemble Construction**: All cell-specific networks form a network ensemble, which is a 3D array (responses × predictor genes × cells). **D. Downstream Analysis**: (D1) NNet captures cellular variation and gene regulation dynamics through co-expression (D2) Non-negative matrix factorisation (NMF) is used to extract principal co-expression patterns, identify transcription factor (TF) modules, and explore their associated regulons. (D3) We adapted the NicheNet framework to integrate gene regulation and signalling interaction databases from OmniPath, constructing integrated prior knowledge networks (PKNs). Annotating cell-specific networks with these PKNs transforms them into contextualised gene regulatory networks (GRNs). In addition, PKNs enable the inference of upstream signalling pathways (USPs) for each cell-specific network by tracing signal transduction paths (receptor–TF–target) inferred based on the contextualised TF–target interactions.

We demonstrate the versatility of NNet through three case studies, with Table 1 summarising the NNet settings and computational costs for each study. First, in Section 2.2, we validate the biological relevance of PCA-approximated co-expression by benchmarking TF activity inferred based on co-expression against that derived from target gene expression alone. Next, in Section 2.3, we analyse the cellular variation in co-expression to capture the dynamic gene regulation occurring during cell differentiation in early haematopoiesis. Finally, in Section 2.4, we integrate cell-specific networks with prior knowledge to investigate signalling interactions within the tumour microenvironment (TME), focusing on those that drive the development of tumour-associated macrophages (TAMs).

**Table 1.** NeighbourNet regression setting and computational cost: We summarise the number of cells (*N*), the number of response genes (*P*), and the number of predictor genes (*Q*) involved in the NNet analysis for each case study. Each NNet analysis thus generates an ensemble of *N* × *P* × *Q* co-expression networks. For NNet analyses in Section 2.3 and Section 2.4, networks were built on a subset of cells (indicated by ‘subsampled’) to represent the full dataset, further reducing computational burden. The **Computational time** column records the runtime of NNet in seconds, broken down into the stages of local gene variance calculation (LVC), regression, and meta-network construction. LVC calculation is an initial part of the co-expression measurement (see Section 4.1) and only needs to be performed once before regression. Adjusting the response genes for subsequent regression steps does not require recalculating local variance. The **Memory usage** column shows the change in memory (in gigabytes) before and after the LVC and regression steps. All the analysis were ran on a RStudio server allocated with four cores of Intel Xeon Gold 6254 @ 3.10GHz CPU. We did not perform parallel computing. Other acronyms: DEGs: differentially expressed genes. PKN: prior knowledge network (see Section 4.2).

| Data | Cells ($N$) | Responses ($P$) | Predictors ($Q$) | Computational time (s) | Memory usage (Gb) |
| --- | --- | --- | --- | --- | --- |
| PBMC3k data<br>Section 2.2 | $N = 2,638$ | $P = 288$<br>CollecTRI TFs | $Q = 4,116$<br>CollecTRI targets | LVC: 34.56<br>Regression: 784.52 | 25.60 |
| Perturb-seq data (d7)<br>Section 2.2<br>[Dixit et al., 2016] | $N = 16,506$ | $P = 10$<br>perturbed TFs | $Q = 3,227$<br>CollecTRI targets | LVC: 872.87<br>Regression: 120.48 | 5.62 |
| Perturb-seq data (d13)<br>Section 2.2<br>[Dixit et al., 2016] | $N = 9,633$ | $P = 10$<br>perturbed TFs | $Q = 3,227$<br>CollecTRI targets | LVC: 305.20<br>Regression: 63.18 | 3.00 |
| Lin- early hematopoietic cell atlas<br>Section 2.3<br>[Pellin et al., 2019] | $N = 1,078$<br>(Subsampled) | $P = 805$<br>PKN TFs | $Q = 4,600$<br>PKN targets | LVC: 33.68<br>Regression: 1045.09<br>Meta-network 482.11 | 34.50 |
| Small cell lung cancer atlas<br>Section 2.4<br>[Chan et al., 2021] | $N = 2,909$<br>(Subsampled) | $P = 28$<br>DEGs | $Q = 900$<br>PKN TFs | LVC: 87.03<br>Regression: 66.61<br>Meta-network 39.11 | 1.23 |

Together, these case studies illustrate NNet’s capability to advance cell-specific GRN inference while revealing insights into complex regulatory processes at a single-cell resolution. NNet, along with its downstream analysis modules, are available as an R package that integrates seamlessly with the widely used Seurat pipeline.

## 2 Results

### 2.1 Overview of NNet

NeighbourNet (NNet) estimates co-expression at the level of individual cells by combining dimensionality reduction with local regression. To measure cell-specific gene co-expression, NNet first uses principal component analysis (PCA) to embed gene expression data into a low-dimensional space (Figure 1A). Within this space, each cell finds its k-nearest neighbours (KNN), and a regression model is fitted to predict the expression of a response gene using the PCs. Co-expression is quantified by examining how strongly each gene embedded within the PCs (i.e. predictor genes) contributes to these local models. By repeating the regression for different response genes, a weighted network is constructed for every cell (Figure 1B–C).

This framework supports both exploratory and hypothesis-driven analyses. To summarise the large ensemble of cell-specific networks, we optimise a non-negative matrix factorisation (NMF) algorithm to identify coherent co-expression patterns across cells and genes, generating soft clusters and enabling the construction of meta-networks that represent these clusters (Figure 1D, top). In parallel, we incorporate prior knowledge by mapping observed co-expression edges to curated regulatory and signalling interaction databases. This step produces context-specific gene regulatory networks (GRNs) with directional information and allows for the inference of upstream signalling pathways (USPs) that connect ligand–receptor interactions to transcriptional responses through contextualised TF-target co-expression (Figure 1D, bottom). The output includes interpretable networks, tools for visual exploration of networks’ regulatory structures, and quantification of active receptor signalling across individual cells.

### 2.2 NNet co-expression between transcription factors (TFs) and targets provides robust evidence for detecting active gene regulation

NNet introduces a novel framework for measuring cell-specific co-expression (Figure 2A1). To evaluate the quality of co-expression estimates obtained from PC regression and to confirm their relevance for gene regulation, we assess whether NNet-derived co-expression between TFs and their targets can improve TF activity inference. If this is the case, it validates NNet’s capability to capture regulatory relationships.

**Figure 2.**
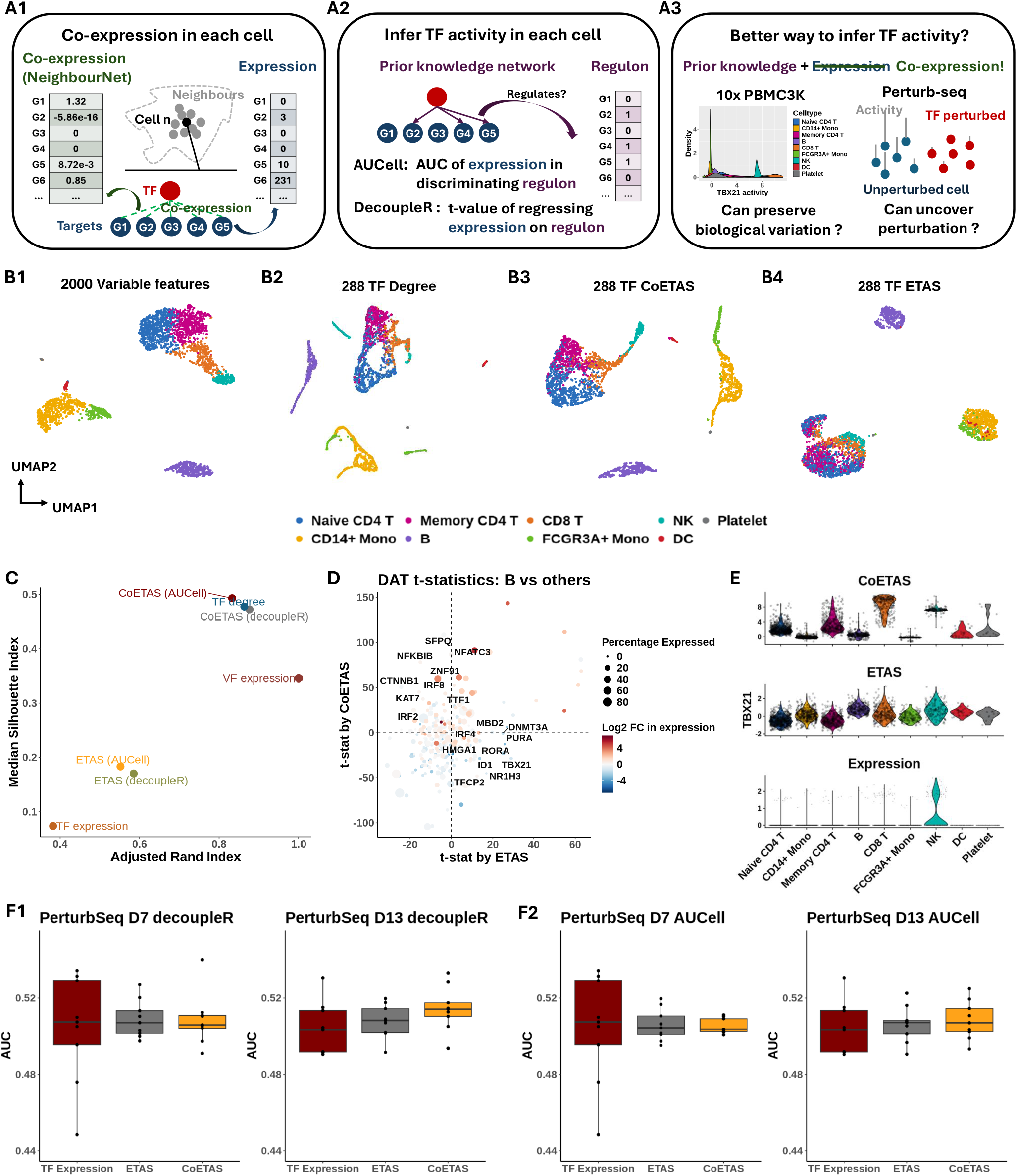
Co-expression between transcriptional factors and targets provides robust evidence to active gene regulation. **(A)** (A1) NNet measures the co-expression between each transcription factor (TF) and its potential targets in each cell’s neighbourhood, yielding a ‘co-expression profile’. We assessed how well these profiles capture TF activity. (A2) Traditional methods (e.g, AUCell and DecoupleR) typically rely on target gene expression to infer TF activity, which can be misleading if high expression does not reflect direct TF regulation. (A3) In contrast, NNet-derived co-expression offers a more reliable measure of true TF activity. AUCell or decoupleR can be applied to co-expression profiles, rather than target expression. We benchmarked all approaches using different measure–method combinations (co-expression vs. expression, with AUCell or decoupleR). **(B)** UMAP visualisation of PBMC 3K data. Cells were embedded with (B1) the 2,000 most variable features (VFs), (B2) node degree for 288 TFs in cell-specific networks, (B3 vs. B4) TF activity inferred from co-expression (CoETAS) vs. expression (ETAS) using decoupleR. CoETAS preserves cell-type boundaries more effectively, resulting in clearer clustering. **(C)** The adjusted Rand and silhouette indexes were used to quantify clustering quality based on TF activity scores. Including the 2,000 VFs and 288 TF expression as controls confirms CoETAS’s superior clustering performance, matching (B3). **(D)** T-tests on decoupleR-derived CoETAS and ETAS identified differentially activated TFs. The x- and y-axes show t-statistics from ETAS and CoETAS, respectively. TFs are coloured by their log-fold change expression in B cells vs. other cell types and sized by the percentage of B cells expressing the TF. Many TFs marked as active by ETAS alone (e.g., TBX21—a T/NK cell regulator) have low expression in B cells, suggesting false positives. **(E)** Comparing TBX21 activity across cell types shows that ETAS (top) mistakenly marks TBX21 as active in B cells, while CoETAS (middle) correctly highlights its known role in T/NK cells. The bottom panel shows TBX21’s log-normalised expression. **(F)** Perturb-seq benchmarking: For 10 TF knockdowns, ETAS and CoETAS were computed to distinguish perturbed cells from controls via area under the curve (AUC). TF expression was served as a baseline. Results with decoupleR (F1) and AUCell (F2) at days 7 and 13 imply that CoETAS outperformed ETAS, especially on day 13.

Traditional methods for inferring TF activity at the individual cell level only assess the expression of known targets of TFs [Badia-i Mompel et al., 2022] (Figure 2A2). However, high expression of target genes may not indicate active TF regulation, which often results in numerous false positives. NNet increases the robustness of cell-specific TF activity inference by generating TF co-expression profiles, thereby enabling existing inference methods to evaluate whether a TF is co-expressed with its targets within the neighbourhood of each cell (Figure 2A3).

We expect that a robust co-expression measure will possess higher regulatory relevance compared to gene expression itself, and consequently yield TF activity scores that accurately reflect true cellular states. To validate this, we conducted a benchmark study comparing TF activity scores derived from expression and co-expression data, abbreviated ETAS (expression-based TF activity scores) and CoETAS (co-expression-based TF activity scores), respectively. This case study underscores the potential of NNet for enhancing the robustness and biological interpretation of TF activity inference.

#### Data and setting

We analysed two datasets. The first was the PBMC 3K dataset from 10x Genomics, often used as a benchmark. The second was a Perturb-seq dataset [Dixit et al., 2016] where cells were exposed to various combinations of TF knockdowns (perturbations) and were sequenced at 7 and 13 days post-perturbation. TF activity scores were calculated using two popular inference approaches: decoupleR and AUCell [Aibar et al., 2017, Badia-i Mompel et al., 2022].

In the PBMC dataset, no ground truth for TF activation exists, we therefore assess how effectively each score captures biological variation. Specifically, we analysed 288 TFs with sufficient target set coverage based on the CollecTRI gene regulation database [Müller-Dott et al., 2023] (Section 4.3). Using NNet, we computed each TF’s co-expression with 4,116 potential target genes (any gene with a known TF regulator in CollecTRI). From these co-expression profiles, we derived CoETAS for each TF and compared them with ETAS generated from target gene expression. This comparison allows us to see which method better preserves cell-type clustering and highlights known cell-type-specific TFs.

The Perturb-seq dataset includes perturbations on 10 TFs, each applied to a different subset of cells. Similar to our approach with the PBMC data, we used NNet to measure co-expression between perturbed TFs and 3,227 target genes. Subsequently, we calculated ETAS and CoETAS for each of the 10 perturbed TFs across all cells. The precise information on TF perturbations in each cell enabled us to directly assess how effectively the inferred TF activity scores distinguish between known TF perturbation statuses.

#### NNet co-expression-based TF activity scores better preserves cell clusters in PBMC data

We annotated the dataset by first performing cell clustering based on the expression of the 2,000 most variable genes, followed by cell type assignment using cluster-specific marker genes (Figure 2B1). Using NNet, we constructed cell-specific co-expression networks, which enabled novel analyses of cellular variation based on network topology, such as connectivity (i.e., node degree). A UMAP embedding of cells based on the connectivity profiles of 288 TFs within these networks largely preserved the separation of cell types (Figure 2B2).

Using both target gene expression and NNet TF-target co-expression, we computed TF activity scores and assessed how well these scores preserve cell type clusters. UMAP embeddings showed that CoETAS (Figure 2B3) better captured known cell types compared to ETAS (Figure 2B4, Supplementary Figure S1A). Quantitative evaluations using the adjusted Rand index and the silhouette index further confirmed that CoETAS produced higher-quality clustering (Figure 2C). As a control, clustering based on TF expression performed worse than TF activity. Additional evaluations using clusters derived from low-rank approximations of TF expression (Supplementary Figure S1B) ruled out the possibility that the advantage of CoETAS stemmed from the PCA regression used to measure co-expression. These results highlight the ability of NNet co-expression in capturing cell-specific regulatory activity.

#### NNet co-expression detects more meaningful cell-type-specific gene regulation in PBMC data

We then assessed the biological relevance of differentially activated TFs (DATs) identified by activity scores in different cell type clusters. We focused on B cells, which formed the most distinct cluster within the PBMC data (Figure 2B). We conducted t-tests on activity scores to extract DATs in B cells. The t-tests on ETAS and CoETAS revealed largely different sets of DATs (Figure 2D). While most DATs identified by both methods were differentially expressed by B cells, several DATs identified solely by ETAS were actually down-regulated or not expressed at all by the B cell population. A key example was TBX21, a master regulator of T and NK cells [Szabo et al., 2000, Knox et al., 2019], which was deemed active in B cells by ETAS. In contrast, CoETAS correctly highlighted the relevance of TBX21 in T cells and NK cells (Figure 2E). Thus, relying solely on target expression for TF activity inference can yield spurious results, whereas NNet co-expression more accurately reflects genuine regulatory activity.

#### NNet co-expression based TF activity scores distinguish perturbation effects more effectively in the Perturb-seq data

We further examined the robustness of CoETAS by analysing TF perturbation experiments of different durations (7 days and 13 days). Activity scores were calculated for each of the 10 perturbed TFs in both perturbed and unperturbed cells. The area under the curve (AUC) was used to evaluate the effectiveness of these scores in distinguishing perturbed (negative class) from unperturbed cells (positive class). The perturbation effect increased with time, as evidenced by higher AUC values for both ETAS and CoETAS (Figure 2F). While CoETAS performed comparably to ETAS on day 7, its superior performance on day 13 suggests that NNet co-expression offers a more accurate detection of TF activation status.

Overall, our findings show that incorporating NNet co-expression significantly enhances the accuracy and biological relevance of TF activity inference, underscoring its potential in effectively capturing gene regulation activities.

### 2.3 NNet cell-specific co-expression networks enable granular profiling of dynamic alternation in gene regulation

Traditional gene regulatory network (GRN) inference methods based on scRNA-seq data typically first cluster cells into groups based on their gene expression and then measure co-expression within these groups [Pratapa et al., 2020]. While effective for describing static and coarse-grained interactions, these approaches struggle to resolve continuously evolving gene regulatory mechanisms, such as transient gene interactions during cell state transitions commonly seen in processes such as cell differentiation.

Using a case study on stem cell biology, we present a novel approach for analysing cellular variation through cell-specific co-expression (Figure 3A1). We propose an efficient matrix factorisation technique to summarise and interpret large-scale co-expression data as meta-networks and meta-genes, revealing overarching co-expression patterns and principal regulatory programs (Figure 3A2). We demonstrate that the NNet framework facilitates accurate identification of pivotal points in cell fate decisions and enhances our understanding of the underlying transcriptional mechanisms that coordinate these decisions.

**Figure 3.**
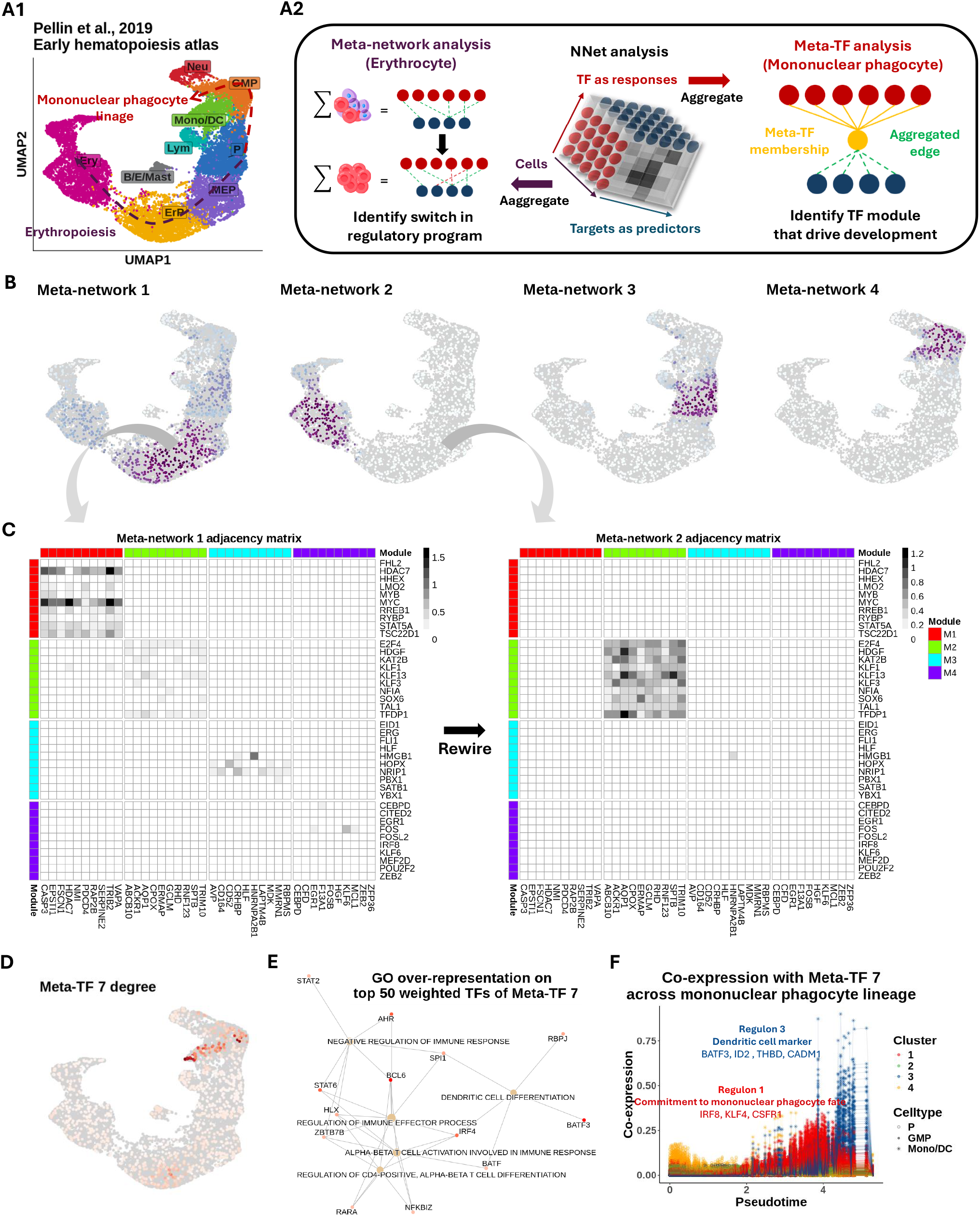
Dynamic co-expression network reveal key transcriptional program alternation during early hematopoiesis. **(A)** UMAP visualisation of bone marrow Lin-cells, depicting the transcriptional landscape of early hematopoiesis [Pellin et al., 2019]. P: progenitor, GMP: granulocyte-monocyte progenitor, MP: mononuclear phagocyte, Neu: neutrophil, Lym: lymphoid cell, Mast: Mast cell, MEP: megakaryocyte–erythroid progenitor, ErP: erythroid progenitor, Ery: erythrocyte. **(B)** Bipartite co-expression networks were constructed for individual cells, linking transcription factors (TFs) to target genes. Non-negative matrix factorisation (NMF) was applied to (1) project the cell dimension into meta-networks and (2) decompose the TF dimension into meta-TFs. Meta-network and meta-TF analyses were used to resolve erythropoiesis (B–C) and mononuclear phagocyte (MP) differentiation (D–F). **(B)** NNet meta-network analysis learns cell weights to soft-cluster and aggregate cells by their co-expression, generating meta-networks. Projecting these cluster weights onto UMAP reveals that the first four meta-networks capture key transcriptional program shifts during myelopoiesis. **(C)** Weighted adjacency matrices of meta-networks 1/2 (early/late erythropoiesis). Top 10 TFs (rows) and targets (columns) from each meta-network reveal rewiring from a stemness-associated state (meta-network 1) to erythrocyte-autonomous programs (meta-network 2), indicating an abrupt transcriptional transition. **(D)** Degree of meta-TF 7 across cells projected onto the UMAP. Meta-TF 7 emerged as the first module associated with the MP lineage, displaying the highest connectivity within GMPs. **(E)** Over-representation analysis of the top 50 TFs that composes meta-TF 7 identifies immune differentiation pathways. TFs linked to the enriched terms are coloured based on their NMF weighting. **(F)** Meta-TF 7’s co-expression with each target gene along pseudotime. Each scatter path represents a target. Targets were clustered into four groups based on their co-expression patterns, with clusters 1 and 3 showing staggered activation waves linked to MP commitment and CD141+ dendritic cell maturation.

#### Data and setting

We used the lineage-negative (Lin-) bone marrow cell scRNA-seq data from [Pellin et al., 2019], which was designed for profiling the early human haematopoietic landscape. With TF and target gene sets acquired from our integrated prior knowledge network (PKN) of gene regulation (described in Section 4.2), NNet analysis was performed to measure co-expression between 805 TFs and 4,600 potential targets.

In the following sections, we first apply meta-network analysis, followed by an examination of cell-specific network structures to characterise transcriptional program alterations during erythropoiesis. We then conduct meta-TF analysis to identify the key TF module that potentially coordinates mononuclear phagocyte (MP) differentiation.

#### NNet meta-network analysis reveals abrupt rewiring of co-expression network during erythrocyte maturation

We identified 20 meta-networks, each characterised by a set of importance score on cells (i.e. soft clustering weights) indicating the cell populations they represent (Supplementary Figure S2A, Section 4.2). These meta-networks captured key transcriptional shifts during hematopoiesis. Specifically, the first four meta-networks delineated the primary myelopoietic lineages towards erythroid and MP fates (Figure 3B). For example, meta-network 1, primarily composed of networks of megakaryocyte–erythroid progenitors (MEPs), was associated with early lineage commitment characterised by HDAC7-driven suppression of MYC that releases erythroid differentiation blockage (Figure 3C) [Jayapal et al., 2010, Delgado and León, 2010, Wang et al., 2020]. In contrast, meta-network 2, which exhibited increased involvement during terminal erythropoiesis, highlighted a shift towards network modules related to haemoglobin synthesis (e.g., KLF1-mediated ABCB10 expression) [Tallack et al., 2010] and erythrocyte membrane protein synthesis (e.g. TAL1-mediated RHD and ERMAP expression) [Kassouf et al., 2010]. Furthermore, the clustering weights of meta-network 2 sharply localised to erythroid progenitors (ErPs) from MEPs, suggesting that erythrocyte maturation may involve abrupt, temporally distinct transcriptional reprogramming. Our results recapitulate a well-established TF regulation mechanism known as the ‘GATA switch’, in which the repression of GATA2 facilitates a surge in GATA1 expression – a crucial step in erythrocyte maturation [Bresnick et al., 2018, Suzuki et al., 2013].

#### NNet network topology of individual cells better characterises temporal dynamics of GATA regulation compared to gene expression

Focusing on the GATA genes (GATA1 and GATA2), we explore how NNet network topology analysis provides an enhanced characterisation of dynamics of transcriptional program compared to conventional gene expression analysis. We calculated the connectivity of GATA genes within cell-specific networks and plotted this connectivity against the diffusion pseudotime of cells (Supplementary Figure 3C1) [Haghverdi et al., 2016]. This analysis showed that GATA2 reached its peak connectivity at the MEP stage, which then sharply declined following the transition to the ErP stage (Supplementary Figure 3C1). As GATA2’s connectivity dropped to zero, GATA1 began to show a significant increase in connectivity throughout maturation, emphasising a critical temporal coordination of GATA2 and GATA1 during erythropoiesis. In contrast, plotting the expression of GATA genes against pseudotime lacks critical details about the dynamics of GATA regulation, such as the peak of activity and the switching point between transcriptional states (Supplementary Figure 3C2).

#### NNet Meta-TF analysis identifies key transcriptional programs that potentially direct CD141+ dendritic cell differentiation

We proceed to illustrate the power of meta-TF analysis, focusing on the identification of key meta-TFs and their regulons associated with MP differentiation, a process distinct from erythropoiesis. Each meta-TF represents a module whose composition is characterised by a set of important scores on TFs. The distinct compositions of different meta-TFs suggest that they represented unique and potentially orthogonal transcriptional programs (Supplementary Figure S2D).

By analysing the connectivity of meta-TFs within each cell (Supplementary Figure S2E), we identified meta-TF 7 as the meta-TF that displayed the strongest connectivity across granulocyte-monocyte progenitors (GMPs) and the MP lineage, implying its potential role in MP differentiation (Figure 3D). To functionally characterise meta-TF 7, we performed a gene ontology (GO) over-representation analysis on its composition. This analysis revealed that meta-TF 7 was closely associated with immune cell differentiation (Figure 3E; analysis details can be found in Section 4.7), particularly the differentiation of dendritic cells (DCs), which belong to the MP lineage. Notably, the DC differentiation term was enriched for BATF3, a key TF that is essential for the development of conventional type 1 DCs (cDC1) [Hildner et al., 2008, Satpathy et al., 2012]. This clinically significant DC subtype plays a critical role in anti-cancer immunity [Böttcher et al., 2018], with extensive research dedicated to developing protocols for its in vitro differentiation [Rosa et al., 2022, Elahi et al., 2022]. We then sought to identify and comprehend potential downstream targets of meta-TF 7 to gain a deeper understanding of its function. Figure 3F illustrates the changes in target gene co-expression with meta-TF 7 during MP differentiation. By clustering targets based on these co-expression values, we identified two major regulons that exhibited staggered waves of activity (Supplementary Figure S2F).

The first wave of activity, represented by regulon 1, included IRF8, KLF4, and CSF1R, which are critical markers indicative of MP fate commitment [Combes et al., 2021, Kurotaki et al., 2014, Tussiwand and Gautier, 2015]. The second wave, represented by regulon 4, featured BATF3 alongside other key markers of cDC1 identity, including ID2, a TF that delineates the cDC1 lineage [Jackson et al., 2011]; THBD, which encodes the CD141 surface marker [Anastasiou et al., 2012]; and CADM1, which is expressed exclusively by cDC1 [Collin and Bigley, 2018]. These findings further support the role of meta-TF 7 in coordinating the transcriptional programs driving cDC1 differentiation. Notably, our analysis of marker expression revealed that cDC1 constituted only a small population at the tip of the MP lineage, making it challenging to identify through clustering analysis alone. This underscores the effectiveness of our meta-TF analysis in disentangling relevant TF-target interactions within rare cell populations.

In conclusion, our analysis of the early hematopoiesis atlas demonstrates NNet’s capability to analyse cellular variation through co-expression, particularly providing insights into the dynamics of gene regulation throughout cell differentiation. By meta-network (meta-TF) analysis, we successfully identified principal co-expression patterns, TF modules and regulons at the individual cell level without the need for cell clustering.

### 2.4 NNet effectively integrates prior knowledge to facilitate cell-specific inference of upstream gene regulation signals

Co-expression between genes does not necessarily imply causal relationships in gene regulation. To enhance the interpretability of co-expression networks, we extend NNet to annotate TF-target co-expression by leveraging prior knowledge of gene regulation and signalling interactions. This approach helps us discern active gene regulation from confounding effects. In addition to identifying regulatory relationships, we infer upstream signalling pathways (USPs), the intracellular signal transduction events that start from receptors and lead to the TF-mediated gene regulation. Moreover, since NNet generates cell-specific networks, the NNet USP inference can also be conducted at the individual cell level, allowing us to further infer signalling dynamics that contribute to cell state transitions.

For our third case study, we analyse scRNA-seq data of small cell lung cancer (SCLC), with a specific focus on its macrophage subset (Figure 4A1). We demonstrate how NNet with prior knowledge annotation comprehensively reveals critical signal interactions within the tumour microenvironment (TME), which may contribute to the development of tumour-associated macrophage (TAM) phenotypes (Figure 4A2).

**Figure 4.**
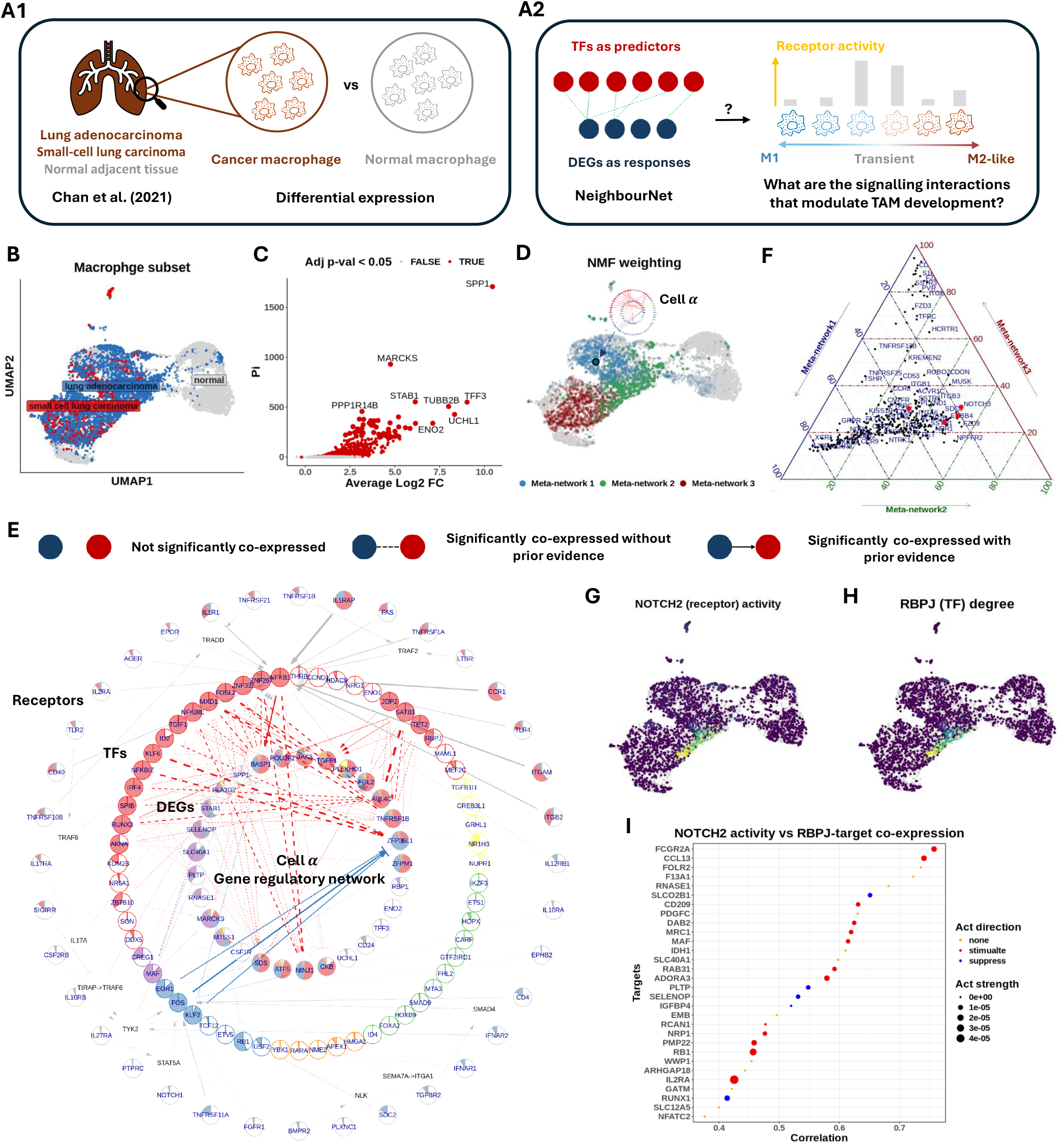
Prior knowledge annotation highlights critical signalling interactions facilitate tumour associated macrophage development. **(A)**(A1) We focused on the macrophage subset of Chan et al. [2021]’s small cell lung cancer scRNA-seq dataset. Using NNet, we aimed to identify the signalling interactions that regulate differentially expressed genes (DEGs) between cancer and normal macrophages, providing insights into how inter- and intra-cellular signaling interactions shape macrophage identity within the tumor microenvironment. (A2) We utilised NNet’s upstream signalling pathway (USP) inference to identify receptors with high activity in driving transcription factors (TF) regulation of DEGs in transient macrophage populations during TAM development. **(B)** UMAP visualisation of macrophages in the dataset. **(C)** Volcano plot of the DE result. Only genes that were up-regulated in cancer are shown. Pi value: negative log10 p-value times log2 fold change. **(D)** Soft-clusters of cells for the first three meta-networks, illustrating a key transition from pro-inflammatory macrophages to TAMs. Cell *α*, with the highest NMF weighting in meta-network 1, is highlighted. **(E)** Annotated co-expression network of cell *α*. The innermost layer contains response (target) genes, encircled by TFs. Receptors occupy the outermost layer, each linked to a TF inferred to mediate that receptor’s influence. If the receptor–TF link is indirect, an extra layer shows the shortest signalling path according to prior knowledge. Section 4.2 provides details. This network captures the canonical NF-*κ*B pathway in pro-inflammatory macrophages. **(F)** Ternary plot of receptor activity scores on the first three meta-networks. NOTCH receptors (NOTCH2/3/4) show peak activity in meta-network 2. **(G)** NOTCH2 activity and **(H)** RBPJ connectivity projected onto the UMAP. RBPJ was the TF whose connectivity was found mostly correlated with NOTCH2 activity. **(I)** Correlation between NOTCH2 activity and different targets’ co-expression with RBPJ. The top targets with the highest correlations are illustrated. NOTCH2 activity is strongly associated with M2-like TAM marker expression mediated by RBPJ.

#### Data and setting

We acquired the SCLC atlas from Chan et al. [2021], which includes scRNA-seq data of both tumour and tumour-adjacent normal tissues (Supplementary Figure S3A). Focusing on the macrophage subset (Figure 4B), we conducted a differential expression (DE) analysis comparing macrophages from tumour and normal samples. Among the top 50 genes up-regulated by tumour macrophages, we identified 29 genes that are targets of known TFs according to our PKN of gene regulation (Figure 4C; the DE analysis is described in Section 4.9). The co-expression of these 29 genes with 900 TFs obtained from the PKN was then measured using NNet. We applied meta-network analysis to recover the major co-expression patterns and subsequently utilised USP inference to investigate how signals are transmitted to the 29 genes through TFs across different macrophage landscapes.

#### NNet annotated co-expression networks capture key signalling pathways defining macrophage interactions within TME

We focused on the top three meta-networks and projected their importance scores onto a UMAP embedding of macrophages (Figure 4D). The importance scores for each meta-network distinctly localised to different macrophage populations, suggesting that each population is governed by a unique transcriptional program. To further explore signalling differences among these populations, we visualised the annotated co-expression networks of the three most representative cells (those with the highest importance in each meta-network), highlighting a spectrum of macrophage identities (Supplementary Figure S3B).

On one end of the spectrum, cell *α*, representing meta-network 1, showed a high level of connection across diverse regulatory modules (Figure 4E). The USP analysis revealed an enrichment of pro-inflammatory signalling in cell *α*, with NFKB1 acting as the central TF that mediated signals from receptors like IL1R, TNFR1, and TLR4. This observation suggests the activation of the well-known NF-*κ*B axis, which strongly supports the M1 pro-inflammatory identity (as opposed to the M2 anti-inflammatory identity) of cell *α* [Yu et al., 2020, Hagemann et al., 2009].

On the other end of the spectrum, cell *γ*, representing meta-network 3, displayed a complete loss of pro-inflammatory signalling (Supplementary Figure S3C1). Instead, it established ENO1-mediated signalling to stimulate SPP1 expression, with SCARB1 as the primary receptor and SRC likely acting as the transducer relaying the signal to ENO1. SPP1+ macrophages represent a distinct subtype of TAM that has been reported to interact with fibroblasts and anti-inflammatory CD8+ T cells, driving tumour growth, metastasis, and immunosuppression [Qi et al., 2022, Bill et al., 2023]. While the roles of SCARB1, SRC, and ENO1 in TAM have been independently reported, their interaction in promoting SPP1 expression presents a potential regulatory mechanism that warrants further investigation [Plebanek et al., 2018, Dwyer et al., 2017, Liang et al., 2023].

In between cell *α* and *γ*, cell *β*, representing meta-network 2, exhibited an intermediate transcriptional state (Supplementary Figure S3C2). We hypothesise that cell *β* and the macrophage population associated with meta-network 2 exist in a highly plastic state, rendering them susceptible to reprogramming and valuable targets for therapeutic treatment. To better understand this transitional state, our next analysis investigates the key signalling interactions and their consequences among these transient macrophages in the TME.

#### NNet receptor activity analysis highlights activation of RBPJ-meditated NOTCH signalling during transitional macrophage state

NNet’s USP inference involves calculating each receptor’s potential activity across target genes for each cell, summarised as activity scores. Leveraging this inference, we explored the receptors that exhibit differential signalling in transient macrophages, represented by cell *β*, compared to M1 macrophages (e.g. cell *α*) and the TAM population (e.g. cell *γ*). Specifically, we summarised receptor activity within each macrophage population by a weighted sum (using importance scores) of cell-specific receptor activities (see Section 2.4). Our comparison revealed that NOTCH2/3/4 receptors were exclusively active in the transient macrophage population associated with meta-network 2 (Figure 4F). Endothelial cells, expressing the NOTCH ligands DLL and JAG, emerged as the principal source of NOTCH signalling (Supplementary Figure S3D). This finding aligns with previous research showing that NOTCH signalling can influence macrophage polarisation towards the M1 or M2 phenotype, although its precise direction remains a debated topic [Palaga et al., 2018, Chen et al., 2023, Xu et al., 2012, Foldi et al., 2016].

To clarify the role of NOTCH in our lung cancer setting, we examined TF connectivity in relation to NOTCH2 activity (Figure 4G). RBPJ emerged as the top TF whose connectivity highly correlated with NOTCH2 activity (Figure 4H), which is consistent with its established role as a key mediator of NOTCH signalling and a frequent target for blocking the pathway [Friedrich et al., 2022]. Our findings partially recapitulate those of Franklin et al. [2014], who reported that RBPJ-mediated NOTCH signalling is crucial for the final differentiation of TAMs from tumour-infiltrating monocytes. However, it remains unclear whether this pathway also governs the macrophage transition from the M1 to TAM phenotype. To address this question, we next investigated the downstream targets of RBPJ to characterise the functional consequences of NOTCH signalling.

#### RBPJ co-expression pinpoints downstream targets of NOTCH signalling and its role in converting macrophages to TAM

We performed a separate NNet analysis to measure RBPJ’s co-expression with 5,011 target genes. By calculating the correlation between NOTCH2 activity (as shown in Figure 4G) and each target’s co-expression with RBPJ across individual cells, we identified a set of genes that were highly associated with NOTCH-RBPJ signalling (Figure 4I). According to prior knowledge, key genes triggered by this signalling include MAF and DAB2, which are important regulators of M2 polarisation [Liu et al., 2020, Marigo et al., 2020], as well as the cell surface receptors MRC1 (CD206) and CD209, which are defining markers for M2-like TAMs [Wang et al., 2024]. While Franklin et al. [2014] suggested that RBPJ-dependent TAMs are MRC1 negative, our findings indicate that RBPJ may induce MRC1 positive TAMs at a transient state. Functionally, we also noted that NOTCH-RBPJ signalling triggered macrophages to secrete chemoattractants such as CCL13, which may recruit immune cells to the TME, thereby modulating the immune landscape (Supplementary Figure S3D).

Taken together, our final case study showcases NNet’s ability to capture signalling interactions that precede gene regulation in individual cells. Through this approach, we identify that NOTCH-RBPJ signalling becomes active at an intermediate macrophage state, positioned between the M1 pro-inflammatory and M2-like TAM phenotypes. In this transitional phase, NOTCH-RBPJ may modulate re-education signals from the TME, facilitating the conversion of M1 macrophages into TAM.

## 3 Discussion

NeighbourNet (NNet) is a novel and highly scalable framework for constructing cell-specific co-expression networks from scRNA-seq data. Based on PC regression, our approach unravels co-expression within the local neighbourhood of individual cells, offering a fresh perspective for understanding regulatory dynamics beyond traditional clustering-based GRN inference methods. A notable strength of NNet lies in its ability to capture networks for thousands of cells with small computational cost. We highlighted the versatility of NNet through three case studies and its utility in facilitating the identification of fine-grained regulatory signals pertinent to specific cell states or transitions.

In the first case study of TF activity inference (Section 2.2), we performed a proof-of-concept analysis on NNet co-expression. By comparing TF activity scores calculated based on TF-target co-expression with those obtained solely from target expression, we demonstrated that co-expression-based scores more accurately reflect TF function and activation status. These findings showed that NNet co-expression can serve as a reliable proxy for gene regulation and highlight that traditional expression-based TF activity inference may yield higher false discovery rates, warranting more cautious interpretation.

In the second case study on early hematopoiesis (Section 2.3), we introduced a novel perspective for analysing cellular variation through co-expression analysis. We showed that analyses traditionally performed in gene expression space, such as dimensionality reduction, clustering, and pseudotime analysis can be effectively adapted to co-expression space. A key insight from this study was that analysing co-expression without relying on predefined clusters substantially improves the ability to uncover detailed regulatory signals in scRNA-seq data. The successful identification of the TF module associated with cDC1 differentiation exemplified this advantage, as traditional cluster-based approaches tend to obscure features of rare cell populations within broader groupings.

In the third case study involving small cell lung cancer (Section 2.4), we demonstrate NNet’s capability to integrate prior regulatory knowledge, ranging from gene regulation to signalling interactions, into the functional characterisation of co-expression networks. Through comprehensive visualisation and quantitative analyses of cell-specific networks, NNet emerges as a pioneering algorithm that resolves fine-grained details of signalling pathways at the individual cell level, addressing a key gap in current computational approaches.

We identified some limitations in our study. First, we did not perform simulation-based benchmarks, which are commonly used to validate new GRN methods for global or cluster-specific networks. This omission is primarily due to the substantial technical challenges of simulating realistic cell-specific networks, which are the focus of our work. Additionally, the lack of competing methods capable of inferring large-scale cell-specific networks across many cells reduces the necessity of such benchmarks. As a result, our validation relied on complex real-world biological datasets. Second, our downstream network analysis focused on basic network properties such as node degree, primarily to enhance interpretability. Incorporating additional metrics, such as clustering coefficients, modularity, or network motifs, could further illuminate the regulatory complexity and dynamic characteristics of cell-specific networks.

Looking ahead, NNet’s framework can be adapted to spatially resolved transcriptomics data, where cell neighbourhoods are defined by spatial proximity rather than gene expression similarity. This extension would enable the study of spatial intercellular communication within tissue-specific microenvironments. Furthermore, by adjusting how data embedding and regression are performed within cell neighbourhoods, NNet can be applied to infer interactions involving molecules beyond transcripts, including ligand–receptor binding (proteomics) and multi-omics relationships. A particularly promising direction is the use of partial least squares (PLS) regression for multi-omics embedding, which could facilitate the integration of diverse data modalities and broaden NNet’s applicability across biological contexts.

## 4 Material & Methods

### 4.1 NeighbourNet: cell-specific co-expression network inference

We develop NeighbourNet (NNet) to infer gene regulatory networks (GRNs) from scRNA-seq data. Unlike traditional methods that rely on coarse-grained cell type clusters to measure gene co-expression, NNet directly analyses co-expression within each cell’s k-nearest neighbourhood (KNN) in the gene expression space. To improve computational efficiency and mitigate noise in scRNA-seq data, we apply principal component analysis (PCA) to embed and denoise gene expression. Co-expression is then measured between PCs and response genes using regression models, and gene-gene co-expression is subsequently recovered from functional dependencies learned between PCs and response genes. In this section, we outline how we build a co-expression network for each cell (cell-specific network) and how we refine these networks based on the KNN graph of cells, a cell-by-cell matrix describing KNN relationships in gene expression. Briefly, the procedure consists of the following steps:

1. Build a KNN-graph based on PCA, describing neighbourhood structure of cells.
2. Fit PC regression within each cell’s neighbourhood to build cell-specific network.
3. Smooth the networks using the KNN-graph to reduce noise.
4. Calculate the significance of co-expression based on the KNN graph constructed.

The technical details are provided in Supplementary Methods S3.3.

#### Build KNN Graph

We perform PCA (centred and scaled) on the gene expression data to extract PCs, with the number of PCs chosen as described in Linderman et al. [2022] (see also Supplementary Methods S3.3). Using these PCs, we reconstruct a low-rank approximation (LRA) of the gene expression data, which serves as the response variable in PC regression. We then construct a weighted KNN graph, where each cell is connected to its neighbours based on distances in the PCA space, following Becht et al. [2019]’s approach (see also Supplementary Methods S3.3). Thirty nearest neighbours of cells are considered as the default. Stronger connections in the KNN graph correspond to shorter distances between cells.

#### PC regression within each cell’s neighbourhood

Within each cell’s neighbourhood, we perform regression using PCs to predict the gene expression of a response gene. Cells are weighted in the regression according to their connection to the master cell of the neighbourhood. Co-expression levels between the response gene and the genes used to compute the PCs (i.e., predictor genes) are recovered from the PC regression model (described later in this section and in Supplementary Methods S3.3). Our framework supports PC regression with various regression methods, with the default being ridge regression with a penalty parameter of 0.5. To reduce noise, the response gene expression is approximated by its LRA reconstruction from the PCs.

Cell-specific networks are constructed by running PC regression with different response genes within each cell’s neighbourhood, thereby measuring co-expression between a set of response and predictor genes. The collection of these cell-specific networks then forms a network ensemble.

#### Permutation feature importance as co-expression measure

Permutation feature importance (PFI) is a straightforward, non-parametric method for assessing the influence of predictors on regression model fitting [Mi et al., 2021]. PFI evaluates the influence of a predictor by permuting its values, making predictions based on the permuted data, and calculating the mean squared difference in prediction outcome across permutations. This methodology can also be applied to PC regression, where we use PFI to measure co-expression between a predictor gene and a response by evaluating how permuting the predictor alters the PCs and subsequently influences the overall prediction in cell neighbourhoods.

However, scRNA-seq data is high-dimensional, making direct permutation on thousands of genes at the neighbourhood level computationally impractical. To address this, we derive an approach for estimating PFI without actual permutation. Our PFI estimation has two main components: the local variance of a predictor gene and the partial derivative of the PC regression model. By combining these components, the estimation can be intuitively interpreted as the predicted squared change in the response given a standard unit change in the predictor expression. A detailed derivation of our PFI estimation and its mathematical formulation can be found in Supplementary Methods S3.3, along with an evaluation of the estimation in Supplementary Results S1.1.

#### Network smoothing with random walk diffusion

Regression within a small neighbourhood of cells can produce unstable estimates in the regression model, rendering cell-specific networks unreliable. We propose to use a random walk diffusion approach, i.e. to smooth the inferred co-expression by propagating co-expression estimates across neighbouring cells with similar expression profiles (see Supplementary Methods S3.3). This propagation is performed iteratively to mimic a random walk on the KNN graph. In each iteration, a cell’s network is updated by averaging them with those of its neighbouring cells, weighted according to the similarity measure derived from the KNN graph. The number of iterations is set to three by default. Consequently, the random walk smooths out random fluctuations in co-expression estimates while preserving important signals that reflect gene-gene interactions.

#### Significance of co-expression

We then evaluate which edges in a cell-specific network represent significant co-expression. To establish a baseline for significance, we assume that a gene is its own perfect predictor. We use the distribution of each response gene’s co-expression with itself (acting as a predictor in the PC regression) across all cells as a reference point.

For each response, we generate a null distribution of co-expression by randomly shuffling its self-co-expression data and then applying random walk diffusion to smooth it over the KNN graph. The random shuffling disrupts the neighbourhood structures in the self-co-expression data, so the smoothing process filters out true signals, leaving behind a baseline that reflects random noise (see Supplementary Methods S3.3 for details). We then compare the co-expression between the response and other predictor genes to this null distribution and calculate probabilistic significance scores (between 0 and 1): if the observed co-expression is substantially larger than what would be expected from the null distribution, the significance score is high. By repeating this process for each response gene, we assess all co-expression estimates in the network ensemble, distinguishing significant edges from those that are not. Users can choose to prune cell-specific networks by defining a significance cutoff for further downstream analysis. By default, the cutoff is set to 0.5 to avoid extensive filtering of edges. In Supplementary Results S1.2, we validated the effectiveness of our proposed pruning approach using an independent dataset from Tian et al. [2019]. Our results demonstrate that the pruning method effectively eliminates co-expression between response genes and random predictors while maintaining a high level of power in retaining significant co-expression.

#### Restrict NNet analysis on subsampled cells to reduce computational demand

To further reduce the computational demands of NNet, both in terms of time and memory usage, and to make it feasible for use on personal computers, we infer cell-specific networks on a representative subset of cells from the entire dataset. To select representative cells, we apply k-means clustering to the PCs, setting the number of clusters to match the desired number of representative cells. For each cluster, we select the cell closest to the cluster centroid as the representative.

NNet runs PC regression and build cell-specific networks on the selected representative cells, leveraging their neighbouring cells from the complete dataset. This yields a smaller network ensemble on which downstream analysis can then be applied. For details on how we perform data smoothing on this network ensemble, refer to Supplementary Methods S3.6.

### 4.2 NNet downstream analysis

Beyond co-expression, NNet supports two major downstream analyses (Figure 1D). We first provide an overview of the two analyses without diving into technical details.

The first is the use of non-negative matrix factorisation (NMF) to derive soft clusters of cells or genes from the ensemble of cell-specific networks. In this approach, each soft cluster of cells is represented as a set of cell weights. Building on these weights, we can construct meta-networks by taking weighted averages of cell-specific networks, thereby capturing overarching co-expression patterns across groups of cells. Similarly, genes can be clustered and aggregated based on their weights that represent gene modules, creating meta-genes and their co-expression profiles across cells. We optimised an existing NMF algorithm to make it scales efficiently on the large ensemble we generate [Shashua and Hazan, 2005].

The second downstream analysis involves annotating observed co-expression using prior knowledge. To achieve this, we constructed integrated prior knowledge networks (PKNs) following the methodology from NicheNet [Browaeys et al., 2020, Müller-Dott et al., 2023], consolidating established regulatory and signalling interaction databases. Significant co-expression edges are assigned directions to represent regulatory effects supported by the PKNs, thereby generating context-specific GRNs. Moreover, we developed a framework to infer upstream signalling pathways (USPs) that drive observed co-expression between TFs and target genes. For each cell in the dataset, its TF-target co-expression network can be used to contextualise PKNs, identifying receptors with high regulatory potential on the targets. Subsequently, we define the USP of a target as the most probable signalling path within the PKNs through which signals from the identified receptors are transduced to the target via contextualised TF-target interactions. Finally, NNet provides comprehensive visualisation tools for exploring these networks and their inferred regulatory and signalling relationships.

#### Embed network ensembles

After constructing the network ensemble, our objective is to identify the most significant co-expression patterns, facilitate soft-clustering of cells based on these patterns. To achieve this, we apply non-negative matrix factorisation (NMF) to decompose the ensemble into meta-networks, as described in Sections 2.3 and 2.4. We opted for NMF due to its enhanced interpretability compared to other factor models such as PCA. Specifically, each cell-specific network is vectorised into a one-dimensional vector, where each entry represents the co-expression between a pair of genes (edges). Collectively, these vectors form a tall matrix, with rows corresponding to edges and columns corresponding to individual cells. Applying NMF to this matrix allows us to extract key co-expression patterns.

However, computing NMF for these large matrices is both time and memory-consuming. Therefore we employ a computationally efficient variant of NMF based on non-negative principal component analysis (nPCA) [Sigg and Buhmann, 2008]. The soft clustering weights used to embed cells in Sections 2.3 and 2.4 correspond to the nPCA loadings learned on cells. We found that applying nPCA directly to the singular value decomposition (SVD) of the tall matrix (see Supplementary Methods S3.4) yields the same results as applying it to the original matrix, allowing us to reduce dimensionality before factorisation. This approach significantly lowers computational costs while preserving key co-expression structures. Additionally, we found that the reduced matrix can be computed directly from the network ensemble without explicitly constructing the full tall matrix, further improving memory efficiency.

A similar nPCA approach is used for meta-transcription factor (meta-TF) analysis (Section 2.3). Instead of retaining the cell dimension, we convert the network ensemble into a tall matrix that preserves the dimension corresponding to TFs. Applying nPCA to this matrix identifies key TF modules, capturing their co-expression profile across the dataset.

Note that, although non-negative tensor factorisation (NTF) [Shashua and Hazan, 2005] could in principle capture these patterns in a single step (treating the ensemble as a 3D tensor without vectorising networks), it was infeasible for our data due to high computational requirements. We therefore used the matrix factorisation approach for practicality and efficiency.

#### Prior knowledge annotation

NNet-derived co-expression are correlative, meaning that it does not necessarily indicate causal regulatory relationships. To improve interpretability, we annotate TF-target co-expression networks with prior knowledge. In addition, observed co-expression provides context for existing regulatory interactions, enabling the inference of upstream signalling pathways (USPs) that may influence target gene expression through TFs.

To enhance the accuracy of our annotations, we first construct integrated prior knowledge networks (PKNs) by combining multiple gene regulation and signalling interaction databases, following the approach of NicheNet [Browaeys et al., 2020]. NicheNet optimises the integration process to construct a PKN that best predicts ligand perturbation outcomes. We use our integrated PKN as the basis for prior knowledge annotation.

#### Construct integrated prior knowledge networks

We performed the NicheNet integration using the R package `Omnipath`, which not only provides a collection of high-quality databases but also offers a framework for users to choose their own databases with prior knowledge confidence levels [Müller-Dott et al., 2023]. See Supplementary Methods S3.5 for details on our database selection process. We obtained two integrated PKNs as directed graphs:

- **TF-target gene regulatory network**: Comprising 1,176 TFs, 6,568 targets, and 41,206 gene regulation interactions.
- **Signalling interaction network**: Incorporating 9,867 genes with 79,242 signalling interactions.

These PKNs were pre-computed and edge-annotated to depict whether they consistently indicate stimulation or inhibition, as aligned in the source databases. Additionally, by executing a personalised PageRank algorithm on the signalling PKN, we pre-computed a matrix reflecting the prior regulatory potential of receptors on TFs (Supplementary Methods S3.5) [Browaeys et al., 2020, Page, 1999].,

- **Regulatory potential matrix**: Comprising the regulatory potential of 415 receptors on 905 TFs, aiding in USP inference.

This matrix will be used to facilitate the USP inference.

#### Calculate receptor activity

To infer USPs for a given TF-target co-expression network, we begin by identifying candidate receptor proteins that are likely to regulate the target genes. This identification hinges on computing a receptor activity score for each candidate receptor, determining how strongly it influences target gene expression through the co-expression network. The core of this approach draws on NicheNet’s ligand activity framework, adapted to our receptor–TF–target setting (see Supplementary Methods S3.5 for more details).

Specifically, we compute a receptor–target activity matrix by multiplying the receptor–TF regulatory potential matrix and the adjacency matrix of the co-expression network (Figure 4A2; see mathematical formulation in Supplementary Methods S3.5). Each entry in this matrix represents the predicted influence of a particular receptor on a target gene, mediated by TFs. By summing these influence values over all targets, we compute a global receptor activity score that reflects the overall strength of a receptor’s regulatory impact on the target gene set. Receptors with high activity scores are considered key upstream regulators and thus prioritised for further exploration.

To incorporate biological knowledge and refine the accuracy of receptor activity estimates, we prune the co-expression network according to three levels of confidence: (i) no pruning, (ii) pruning statistically insignificant edges, and (iii) pruning edges that are either insignificant or unsupported by prior knowledge. We consider an edge to be ‘supported’ if the corresponding TF can transmit its signal to the target within a defined number of steps (2 steps by default) on the gene regulation PKN. The default high-confidence setting (iii) retains only those edges backed by robust co-expression evidence and existing regulatory interactions in the PKN. This pruning strategy enhances the accuracy of our inferred receptor activity scores and ultimately improves the reliability of the identified USPs.

#### Reconstruct upstream signalling pathways (USPs)

After identifying highly active receptors, we next reconstruct their potential signalling routes to the target genes. To do this, we focus on the subset of TFs through which each receptor most plausibly relays regulatory signals, incorporating both co-expression and prior knowledge evidence (details in Supplementary Methods S3.5). In essence, those TFs should strongly co-express with the target genes while being supported by prior knowledge of gene regulation and exhibit a high regulatory potential from the receptor.

With TFs selected, we employ the `igraph::shortest_paths` function from the R package `igraph` (Version 2.0.1.1) to discover minimal signalling routes within the signalling interaction PKN. For each high-activity receptor, the algorithm identifies the shortest path(s) that connect the receptor to its relevant TFs. These paths represent the most direct series of known signalling interactions linking receptor activation to TF modulation.

By integrating these receptor–TF routes with the TF–target interaction contextualised by co-expression, we reconstruct candidate USPs for each receptor–TF–target triplet. This final set of pathways illustrates the chain of biochemical events through which signals travel from an upstream receptor, pass through intermediate signalling components, ultimately reach the TF, and thereby regulate target gene expression.

#### Visualise prior knowledge enriched networks

We visualise prior-knowledge-enriched TF-target co-expression networks using a hierarchical structure, as shown in Figure 4E.

1. **Target genes**: positioned at the inner most layer.
2. **TFs**: encircle targets, and are grouped by expression-based clustering (node color), assigned significance probabilities (pie charts: % fill = likelihood of co-expression with targets).
3. **USPs**: The outermost layer lays the most active receptors, each connected to a TF that likely mediates the receptor’s influence on the targets via USP inference. Receptors are coloured to match TFs, with the percentage of color filling represents the expression level. When a receptor and a TF lack a direct link, USP inference puts an extra layer between them shows their shortest signalling path.

Edges indicate evidence of interaction: none (no edge), significant co-expression only (dashed), or significant co-expression supported by prior evidence (solid, with arrowhead for activation or barhead for repression).

### 4.3 Data for NNet analysis

We provide an overview of the public scRNA-seq datasets utilised in our case study and detail the pre-processing steps performed prior to NNet analysis. All data were pre-processed and analysed based on a `Seurat` object in R (`Seurat` package version: 5.0.3). Unless specifically noted, NNet was configured using the default tuning parameters as described in the methodology section below.

#### The 10x PBMC3k data

We obtained a widely recognised dataset of human peripheral blood mononuclear cells (PBMC), commonly used for benchmarking cell clustering methods, from the Seurat PBMC3k tutorial at https://satijalab.org/seurat/articles/pbmc3k_tutorial (Last retrieved: Oct 27th, 2024) [Butler et al., 2018]. Following the code outlined in the vignette, we pre-processed the data, performed dimension reduction using PCA and UMAP, and conducted clustering. The resulting data is log-normalised, containing 2,638 cells and 13,714 genes.

We treated TFs as responses and target genes as predictors in our NNet analysis. Since our TF activity inference (detailed in 4.4) relies on gene regulation data from the CollecTRI database [Müller-Dott et al., 2023], we extracted the relevant TF and target gene sets from CollecTRI. Among the 13,714 genes in the dataset, 731 TFs and 4,116 target genes were identified. To ensure meaningful inference of TF activity, we further filtered the 731 TFs to include only those with at least 10 associated targets according to CollecTRI prior knowledge. This resulted in 288 TFs for NNet analysis on all the cells, maintaining 4,116 target genes as predictors.

#### The Perturb-seq data

The data was originally generated by Dixit et al. [2016], who established the scRNA-seq protocol for pooled CRISPR perturbation on multiple loci of individual cells to discern genetic interactions. We downloaded the cleaned version of the data produced from the repository of Holland et al. [2020] (https://zenodo.org/records/3564179), who conducted a systematic benchmark of TF activity inference methods using this data. The data comprises 10,000 genes and includes a total of 26,139 cells, each of which could be perturbed with a random combination of 10 different TFs (EGR1, NR2C2, E2F4, CREB1, YY1, GABPA, IRF1, ELK1, ELF1, ETS1), or perturbed inter-genetically as negative controls. The cells were sequenced at two time points: 16,506 cells at 7 days and 9,633 cells at 13 days post-perturbation. We did not perform additional quality control to filter out genes or cells. The data was then log-normalised using the Seurat function `Seurat::NormalizeData` with its default tuning.

The NNet analysis was performed on all the cells using the 10 perturbed TFs as responses. We applied the same procedure as in the PBMC data to select 3,227 targets in CollecTRI as predictors.

#### The early hematopoiesis atlas

The data generated by Pellin et al. [2019] profiles an early human hematopoietic landscape of bone marrow mononuclear cells that lack mature lineage markers (Lin-). We obtained the raw data from the GEO accession GSE117498 and downloaded the four Lin-samples. These samples were merged into a combined data comprising 24,720 genes and 15,397 cells. Without performing additional quality control to filter out genes or cells, we applied the Seurat pipeline to process the data. We log-normalised the data using the default `Seurat::NormalizeData` method. Subsequently, we conducted PCA (centred and scaled) and utilised the top 10 PCs to generate a UMAP projection. From the UMAP, we did not observe technical differences between cells from different samples, hence data integration for batch effect correction was omitted. For cell clustering, we employed `Seurat::Findneighbours` using the top 10 PCs, followed by `Seurat::FindClusters` with the resolution parameter set to 0.4, resulting in nine distinct clusters. We annotated cell clusters according to the marker genes provided by Pellin et al. [2019].

The NNet analysis was conducted on a subset of 1,078 cells, using 805 TFs as responses and 4,600 targets as predictors. These TF and target gene sets were collected from our PKN with the criterion that they should be expressed by at least 20 cells.

#### The small cell lung cancer atlas

The dataset was generated by Chan et al. [2021], who aimed to profile the heterogeneity of small cell lung cancer (SCLC) and its associated microenvironment across different lung cancer subtypes, with a specific focus on identifying and characterising macrophage subpopulations linked to the presence of a metastatic SCLC subtype. Additionally, they collected and sequenced specimens from lung adenocarcinoma (LUAD) and normal lung tissues for comparative analysis. We downloaded the processed data from the CELLxGENE data portal, which comprises 22,397 genes and 147,137 cells pooled from 42 donors. We retained 19,558 genes that have HUGO symbols for easier interpretation and did not perform additional quality control to filter out cells from the processed data. Utilising the cell type annotations provided by CELLxGENE, we extracted 9,251 macrophages, on which we then performed PCA (centred and scaled) and UMAP (with 10 PCs). We did not observe significant batch effects; therefore, data integration was not performed.

We conducted two separate NNet analyses on the same set of 2,909 cells that were sub-sampled (Section 4.1): one aimed at identifying signalling interactions associated with pro-tumorigenic macrophage development, and another focused on determining the downstream targets of NOTCH-RBPJ signalling. For the first analysis, response genes were identified through a differential expression (DE) analysis comparing macrophages from tumour and normal samples (Section 4.9). From the top 50 differentially expressed genes (DEGs) of tumour macrophages, we selected 28 genes with known TF regulators according to our PKN to serve as responses for NNet analysis. In the second analysis, RBPJ was set as the NNet response. When selecting predictors, we only considered genes that were expressed by at least 20 macrophages. As a result, 900 TFs and 5,011 targets (not specific to RBPJ) in the PKN were selected as predictors for the first and second analyses respectively.

### 4.4 AUCell and decoupleR analysis for TF activity inference

The R package `decoupleR` (Version 2.9.7) provides an ensemble of computational approaches for TF activity inference based on gene expression data [Badia-i Mompel et al., 2022]. This ensemble includes the AUCell method and the package’s new approach that relies on univariate linear regression, referred to as the decoupleR method in our paper [Aibar et al., 2017]. Both AUCell and the decoupleR method infer TF activity by evaluating how well a cell’s gene expression can predict the known target gene set of a TF. AUCell, executed by the function `decoupleR::run_aucell`, calculates the area under the curve (AUC) score for the sorted expression of a cell in distinguishing the target gene set. In contrast, the decoupleR method, executed by `decoupleR::run_ulm`, performs a regression between gene expression and a dummy vector representing the target gene set, calculating TF activity as the t-value of the regression. Both functions take a gene expression matrix and a prior knowledge GRN as inputs, iterate AUC and t-value calculations on each cell for each TF, eventually generating a TF-by-cell matrix of TF activity. The prior knowledge GRN we used was generated by `decoupleR::get_collectri` based on the CollecTRI database, which was in a specific data format required by the decoupleR functions.

TF activity inference based on TF-target co-expression was also performed using the decoupleR package. The only difference was that, for each TF, we measured its co-expression with targets across cells, generating a TF-specific co-expression profile on which activity inference was then performed.

### 4.5 Evaluate the effectiveness of TF activity in distinguishing perturbed cells using area under the curve

For the Perturb-Seq data, we used AUCell and decoupleR to measure the activity of 10 TFs that were perturbed. For each of the 10 TFs, we compared the activity scores between two groups of cells: those perturbed exclusively on that TF and those that were not perturbed on any TF. The area under the curve (AUC) was calculated to assess how well the activity score of the perturbed TF could distinguish the unperturbed cells.

In both the data sequenced at 7 and 13 days post-perturbation, we observed two significant batch clusters: 3,701 and 12,805 cells at day 7, and 2,600 and 7,033 cells at day 13. The AUC scores shown in the main result (Section 2.2) were calculated based on the larger batch cluster for each time point. AUC evaluations for the smaller batch clusters are provided in Supplementary Figure S1. AUC was calculated using the R package `pROC` (Version 1.18.0) with the function `pROC::roc`.

### 4.6 Embed and cluster PBMC3K data by TF activity

On the PBMC data, we calculated TF activity for 288 TFs across 2,638 cells based on different activity inference methods (AUCell and decoupleR) applied to different measurements (expression and co-expression). The resulting TF activity matrices (Section 4.4) were then embedded using PCA (centred and scaled) followed by UMAP (using 10 PCs).

We employed the adjusted Rand index (ARI) and median silhouette index (MSI) to quantitatively assess how well the activity scores of 288 TFs recapitulate the cell clusters we learnt from the expression of the 2000 most variable features [Rand, 1971, Rousseeuw, 1987]. MSI was calculated based on the distances between cells in the 10 PC space of activity scores, evaluating whether cells within the same gene expression cluster are also grouped together according to their TF activity profiles. ARI is a metric that evaluates the agreement between two different clustering schemes. We utilised ARI to compare clusters generated from activity scores with those derived from gene expression data. For the clustering based on activity scores, we employed Seurat’s clustering pipeline: `Seurat::Findneighbours` was run on the 10 PCs of activity scores, followed by `Seurat::FindClusters`, which was tuned to generate exactly 9 clusters, matching the number of clusters obtained from the expression data.

### 4.7 Over-representation analysis on meta-TF weighting vectors

We performed an over-representation analysis on the NMF weighting vector of meta-TF 7, the first meta-TF that showed strong connections with target genes in the mononuclear-phagocyte lineage. Using the R package `clusterProfiler` (version 4.11.1) and its `clusterProfiler::enrichGO` function, we identified over-represented gene ontology (GO) terms in biological processes among the top 50 weighted TFs of meta-TF 7 [Yu et al., 2012]. A false discovery rate threshold of 0.05 was set to determine significant GO terms, and the background gene set for the analysis consisted of the 805 TFs used in the NMF embedding.

The resulting GO terms were ranked according to their pi-values, calculated as the product of their fold changes and their negative log10 p-values [Xiao et al., 2014]. To address redundancy among highly similar GO terms, we applied the `clusterProfiler::simplify` function, selecting the GO terms with the highest pi-value to represent each group of similar terms. Finally, the top five non-redundant GO terms with the highest pi-values were illustrated.

### 4.8 Diffusion pseudotime inference

Using the R package `destiny` (Version 3.10.0), we performed diffusion pseudotime (DPT) inference on the early hematopoiesis atlas to reconstruct the temporal ordering of cells during differentiation [Haghverdi et al., 2015, 2016]. Briefly, DPT for each cell was derived from its distance to a user-selected root cell, representing the earliest differentiation stage, within a diffusion map (DM) embedding of the data. To create the DM, we applied `destiny::DiffusionMap` to the 10 PCs of the 2,000 most variable genes computed previously (Section 4.3). The resulting DM served as the input for `destiny::DPT` to infer DPT. We selected the root cell as the one with the highest CD34 expression, a marker for hematopoietic stem cells.

### 4.9 Differential expression analysis on macrophages within lung tumour

Macrophages from SCLC and LUAD samples were grouped together for comparison with normal macrophages, as differences between cancer subtypes were not of interest. The DE analysis was performed using the Seurat function `Seurat::FindMarkers`, selecting MAST as the DE method and retaining DEGs that were expressed in at least 10 percent of the tumour macrophages [Finak et al., 2015]. The top 50 DEGs of the tumour macrophages were then selected according to their pi-values [Xiao et al., 2014].

## 5 Code availability

Reproducible code and the `NeighbourNet` R package is available at github.com/meiosis97/NeighbourNet

## 6 Declaration

## S1 Supplementary Results

### S1.1 Evaluate linear approximation of permutation feature importance as the co-expression measure

NNet’s gene co-expression measure is derived from permutation feature importance (PFI). In the standard procedure, after fitting a PC regression model for a response gene within a cell’s neighbourhood, we permute the expression values of a predictor gene within the neighbourhood while keeping all other predictor genes unchanged. The permuted data is then projected onto the global PCA loading, and the resulting PCs are fed into the fitted regression model to predict the response. PFI for the permuted predictor gene is calculated as the expected squared difference between the prediction made before and after permutation. However, directly permuting each predictor gene and making repeated predictions for thousands of cell neighbourhoods is computationally prohibitive. To address this, we derived a linear approximation of PFI that can be easily calculated, bypasses the permutation-and-prediction procedure (Supplementary Methods S3.3.2). Here, we evaluate how well this approximation performs compared to the PFI scores calculated using actual permutation.

We utilised a human lung adenocarcinoma cell line (HCC827, H1975, A549, H838 and H2228) dataset, containing 3,918 cells, for evaluating the goodness of the approximation [Tian et al., 2019]. We downloaded the processed data from https://github.com/LuyiTian/sc_mixology/tree/master/data (Last retrieved: Dec 12th, 2024). PCA was conducted on the 2,000 most variable features (VFs) to generate global PCs. We then randomly selected cells and their neighbouring cells to perform PC regression using different regression methods: ridge regression, random forest (RF), and support vector machine (SVM) with a radial kernel. For each randomly sampled cell, a gene from the 2,000 VFs was randomly chosen as the response, and another gene was selected to measure its PFI and approximated PFI in predicting the response. This procedure was repeated 1,000 times for diferent cells and genes, with PFI scores calculated using both actual permutation and the approximation for each regression method. We then evaluated the correlation between the approximated and actual PFI scores across the iterations.

The results demonstrate that the approximated PFI aligns perfectly with the actual PFI for linear regression methods such as ridge regression, showing near-perfect correlations. For SVM with a radial kernel, the approximation also performed exceptionally well, with correlations exceeding 0.9, indicating strong concordance between the approximated and actual PFI. However, the approximation was less effective for RF, yielding weaker correlations. This discrepancy may be due to the smaller sample size within each neighborhood, which exacerbates the discontinuous nature of RF predictions. Since the PFI approximation relies on estimating partial derivatives, discontinuities in RF predictions likely introduce errors, reducing the accuracy of the approximated PFI.

Overall, our linear approximation of PFI provides a highly efficient and accurate alternative to permutation-based PFI for linear models and SVMs, significantly reducing computational burden while maintaining reliability. While it is less effective for RF, this limitation underscores the dependency of the approximation’s accuracy on the smoothness of the underlying regression model’s predictions.

### S1.2 Sensitivity and specificity analysis of NNet co-expression pruning based on significance measures

We performed a sensitivity (false positives rate) and specificity (true negative rate) analysis to evaluate the effectiveness of NNet pruning in distinguishing meaningful from irrelevant gene associations using the proposed significance measure on co-expression (Supplementary Methods S3.3.5). A key feature of PC regression, leveraged in this analysis, is its ability to measure co-expression between response genes and themselves (self-co-expression) when these response genes are also used to embed PCs. Since a gene should inherently exhibit a significant self-co-expression, this provides a natural benchmark for assessing the pruning process. Specifically, we examined whether NNet pruning successfully retains these self-co-expression, which represent true positive associations, while filtering out irrelevant or non-significant co-expression, minimising false discoveries.

We used the human lung adenocarcinoma cell line dataset described in Section S1.1 for this analysis, focusing again on the 2,000 most VFs. To evaluate specificity, we first extracted the expression data of these VFs and generated a permuted dataset as a negative control, where each VF was permuted independently. The original and the permuted dataset were concatenated, forming a single dataset with 4,000 features (2,000 original VFs and 2,000 permuted VFs). PCs were computed on this concatenated dataset, and NNet analysis was performed using each un-permuted VF as the response. This allowed us to measure and compare each VF’s self-co-expression and the co-expression with its permuted counterpart across cells. A pruning selection was considered a true positive if a VF’s self-co-expression was identified as significant in a cell, and a false positive if the co-expression with the VF’s permuted counterpart was identified as significant. Using this framework, we calculated the sensitivity and specificity for each VFs.

For each VF, we performed the analysis separately in cell populations where the VF was expressed and not expressed. This stratification accounts for the expectation that a VF should be expressed to exhibit significant self-co-expression. The results are summarised in Supplementary Figure S5A. Under NNet’s default pruning settings, cells expressing the VF achieved a sensitivity of approximately 0.6 in selecting self-co-expression. As expected, cells not expressing the VF showed lower sensitivity. In terms of specificity, NNet effectively rejected co-expression with permuted counterparts, achieving near-perfect specificity across both cell populations. These results indicate that NNet’s default pruning settings are conservative, favouring the rejection of false positives.

Despite this overall success, we identified VFs where NNet exhibited very low sensitivity. To investigate, we examined the distribution of importance scores representing self-co-expression and co-expression with permuted counterparts, along with the null importance distribution proposed in Section S3.3.5, which is used to determine significance thresholds. For most VFs, the null distribution clearly separated importance scores of themselves from those of their permuted counterpart, effectively supporting accurate pruning (Supplementary Figure S5B). However, in scenarios where the VF was ubiquitously expressed across all cells and all self-co-expression should be considered significant, the null distribution forced a classification of significant and non-significant (Supplementary Figure S5C), highlighting a limitation of the current approach in handling such cases. One potential improvement is to construct the null distribution of co-expression based on the most variable co-expression between a predictor gene and the response, rather than using the response’s self-co-expression. The rationale behind this approach is that the most variable co-expression values are more likely to encompass a broader range of the co-expression distribution, thereby creating a null distribution that provides a clearer distinction between significant and insignificant co-expression.

## S2 Supplementary Figures

**Figure S1.**
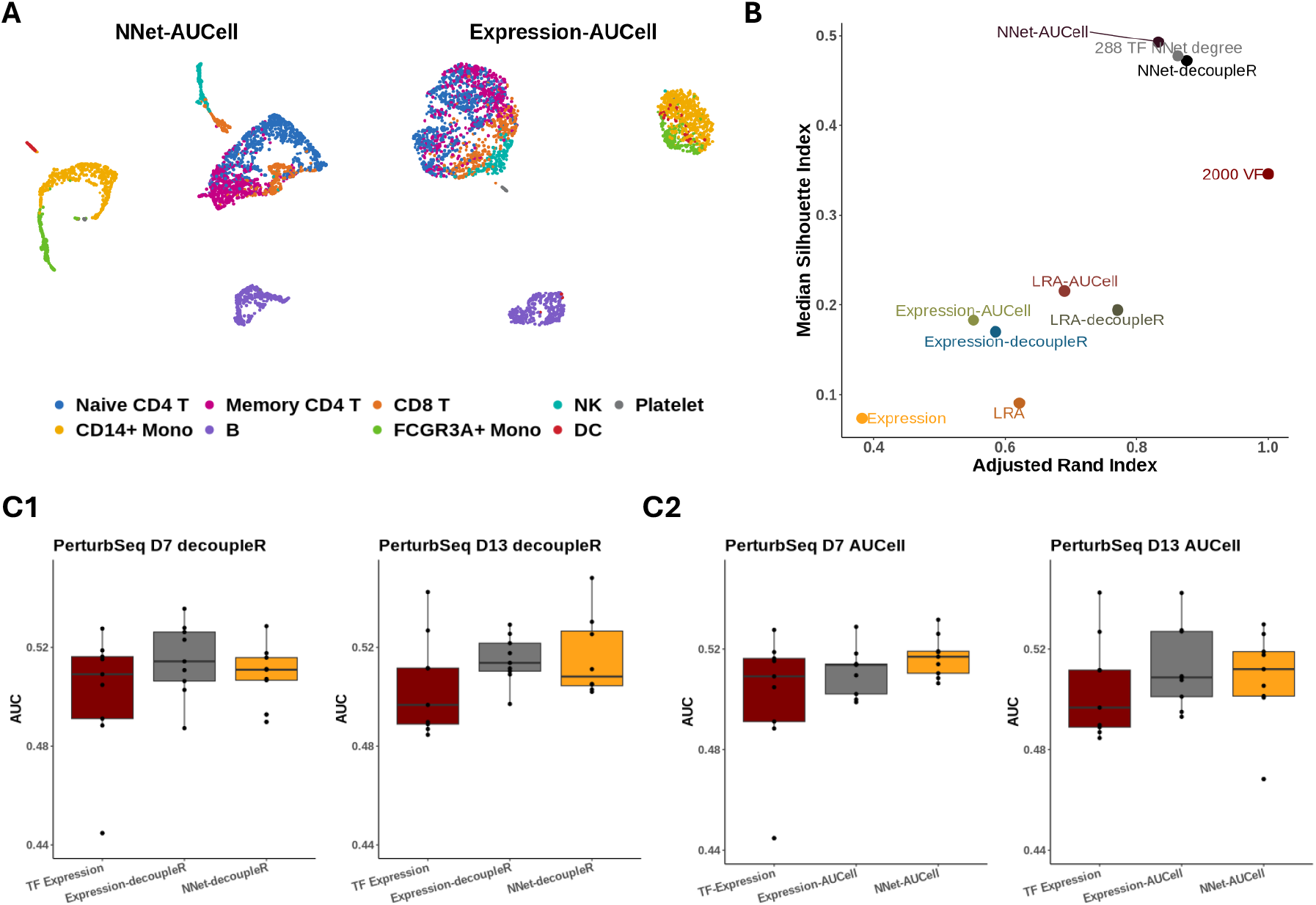
TF activity case study (Section 4.4). **(A)** Similar to Figure 2B, UMAP visualisation of 10x Genomics PBMC 3K data, embedded based on the TF activity scores inferred by AUCell. **(B)** An extension of Figure 2C, comparing clustering schemes of the PBMC 3K data using 10 principal components (PCs) derived from various TF activity measures. The additional measures include TF activity scores based on low-rank approximation (LRA) of gene expression using AUCell (LRA-AUCell) and decoupleR (LRA-decoupleR). The LRA of 288 TF expressions is included as a control. Clustering evaluation, labeled as ‘200 TF NNet degree’, was also performed on the PCs that were used to compute the UMAP in (A). **(C1)** and **(C2)** are similar to Figure 2F1 and 2F2, benchmarking the effectively of TF activity in distinguishing perturbed cells of the Perturb-seq data. The area under the curve (AUC) scores in (C) and Figure 2F1 were calculated across batch clusters of different sizes. Specifically, (C) shows the AUC results for the smaller batch cluster, while Figure 2F1 presents the results for the larger batch cluster.

**Figure S2.**
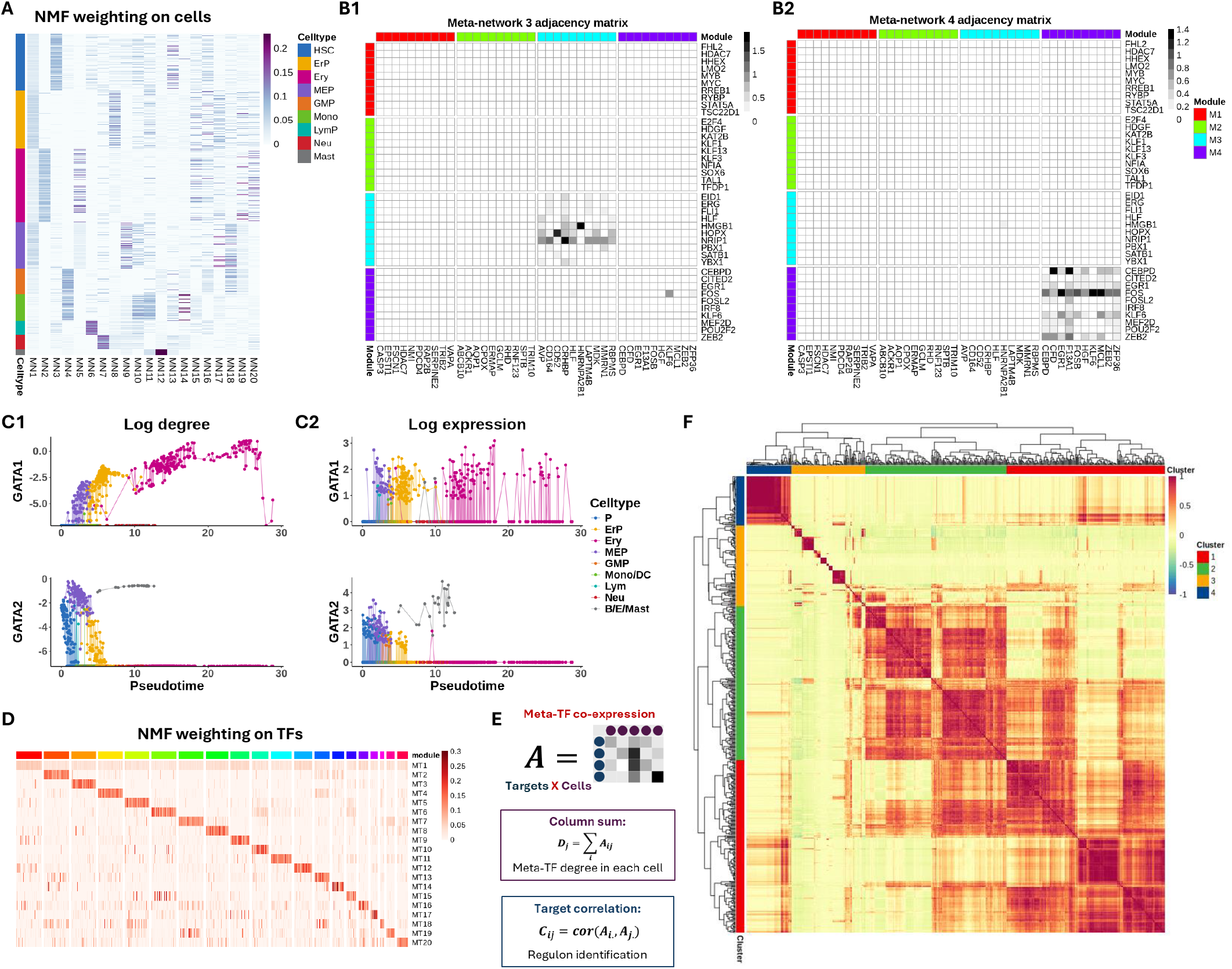
Meta-network case study on early hematopoiesis (Section 2.3). **(A)** Heatmap of cell weights derived from NMF, which were used to embed the cell dimension of the network ensemble. Each column represents one of the 20 weighting vectors used to construct the 20 meta-networks. The weight of a cell indicates its relevance to the corresponding meta-network. **(B)** Weighted adjacency matrix representation of meta-network 3 and 4. The top four most connected TFs and their targets were selected within each of the first four meta-networks, represented as rows and columns, respectively. Genes are grouped by the meta-network from which they were extracted. **(C)** Continuous tracking of GATA gene (GATA1 and GATA2) regulatory dynamics during erythropoiesis. (E1) tracks changes in connectivity (log node degree) for GATA genes within cell-specific networks during erythropoiesis. Compared to the gene expression patterns shown in (E2), connectivity provides a more precise representation of the regulatory dynamics of these TFs, highlighting a distinct switching behaviour between GATA2 and GATA1. **(D)** Similar to A, but the rows of this heatmap illustrates the 20 NMF weighting vectors that embed TFs in the network ensemble. Each weighting vector represents a TF module that contributes to constructing a meta-TF, with the weights indicating the contribution of individual TFs. **(E)** Individual TF co-expression with targets across cells (represented as a target-by-cell matrix) can be aggregated using the weighting vector of an identified meta-TF. This aggregation represents the co-expression between the meta-TF and its targets in each cell. meta-TF co-expression can be analysed by (1) summing target co-expression within each cell to quantify meta-TF activity, and (2) calculating correlations between targets to identify the meta-TF regulon. Here, we focus on TF module 7, which was found to be differentially connected in the mononuclear-phagocyte lineage according to our first analysis proposed. **(F)** Target correlation calculated based on their co-expression with meta-TF 7. From hierarchical clustering we identified four major regulons.

**Figure S3.**
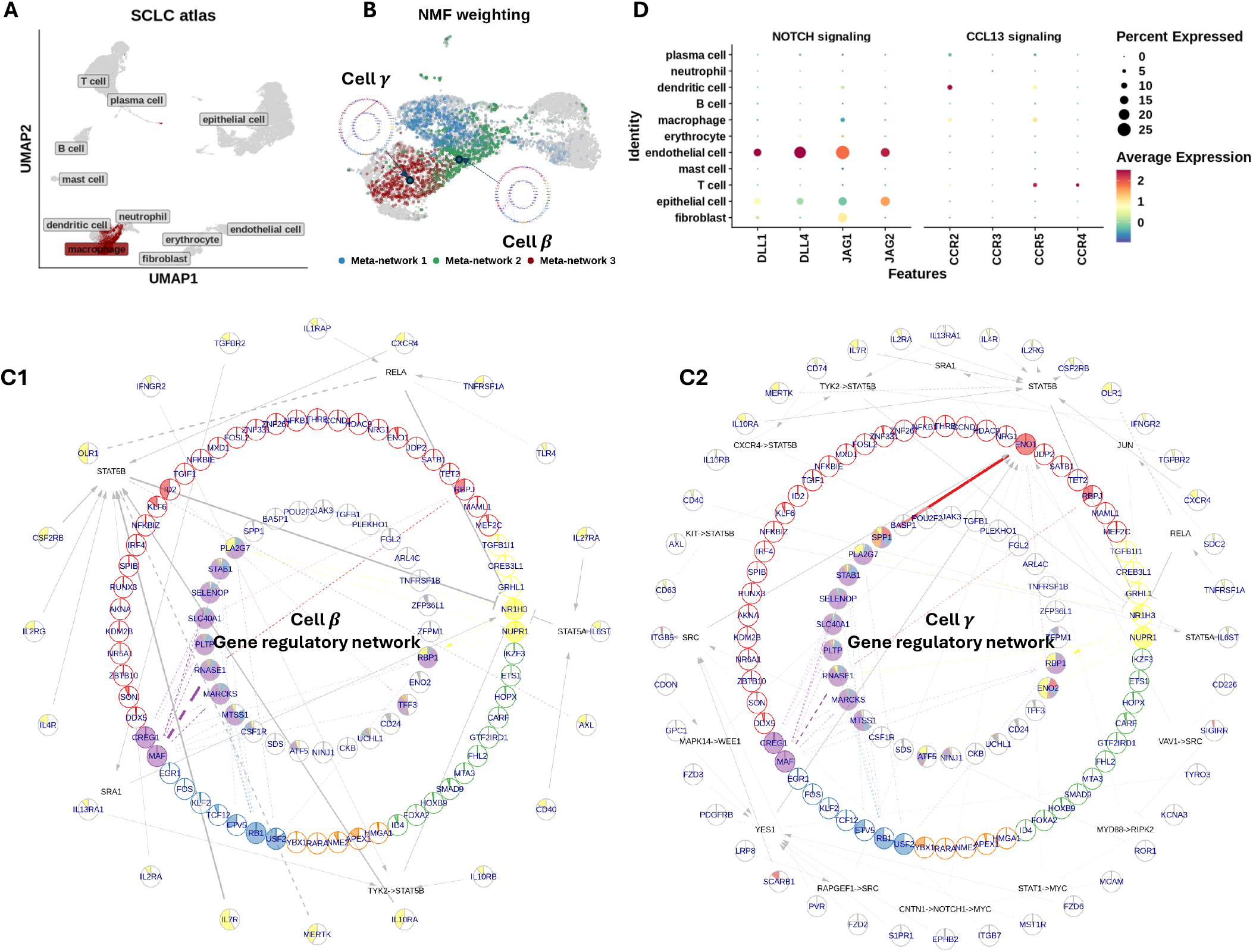
Prior knowledge annotation case study on tumour associated macrophage (Section 2.4). **(A)** UMAP visualisation of the small cell lung cancer scRNA-seq data of Chan et al. [2021]. The UMAP embedding was pre-computed and stored by the CELLxGENE scRNA-seq data portal. **(B)** Continue of panel (D) from 4. Cell *β* and cell *γ*, which has the highest NMF weighting in meta-network 2 and 3, are highlighted respectively. **(C)** Prior knowledge annotation of the co-expression network of cell *β* (C1) and cell *γ* (C2). Cell *γ* displays a dual identity involved in a NR1H3 mediated pro-inflammatory response, undergoing suppression directed by anti-inflammatory signals from IL4R and IL13R. Cell *γ* exhibits decreased level of NR1H3 connection and an establishment of a connection between ENO1 and SPP1, characteristic of an SPP1+ pro-tumourigenic macrophage. According to our upper-stream signalling pathway (USP) inference, a top USP candidate for ENO1-mediated SPP1 signalling in cell *γ* is SCARB1, with its signal likely transduced via SRC. **(D)** The dot plot illustrates the expression levels of key ligands involved in the NOTCH signalling pathway alongside the expression of the CCL13 receptor across various cell populations. Endothelial cells could be the major source of NOTCH signalling that contributes to tumour associated macrophage development. As a response, macrophages of intermediate states release CCL13 that predominantly targeting immune cells.

**Figure S4.**
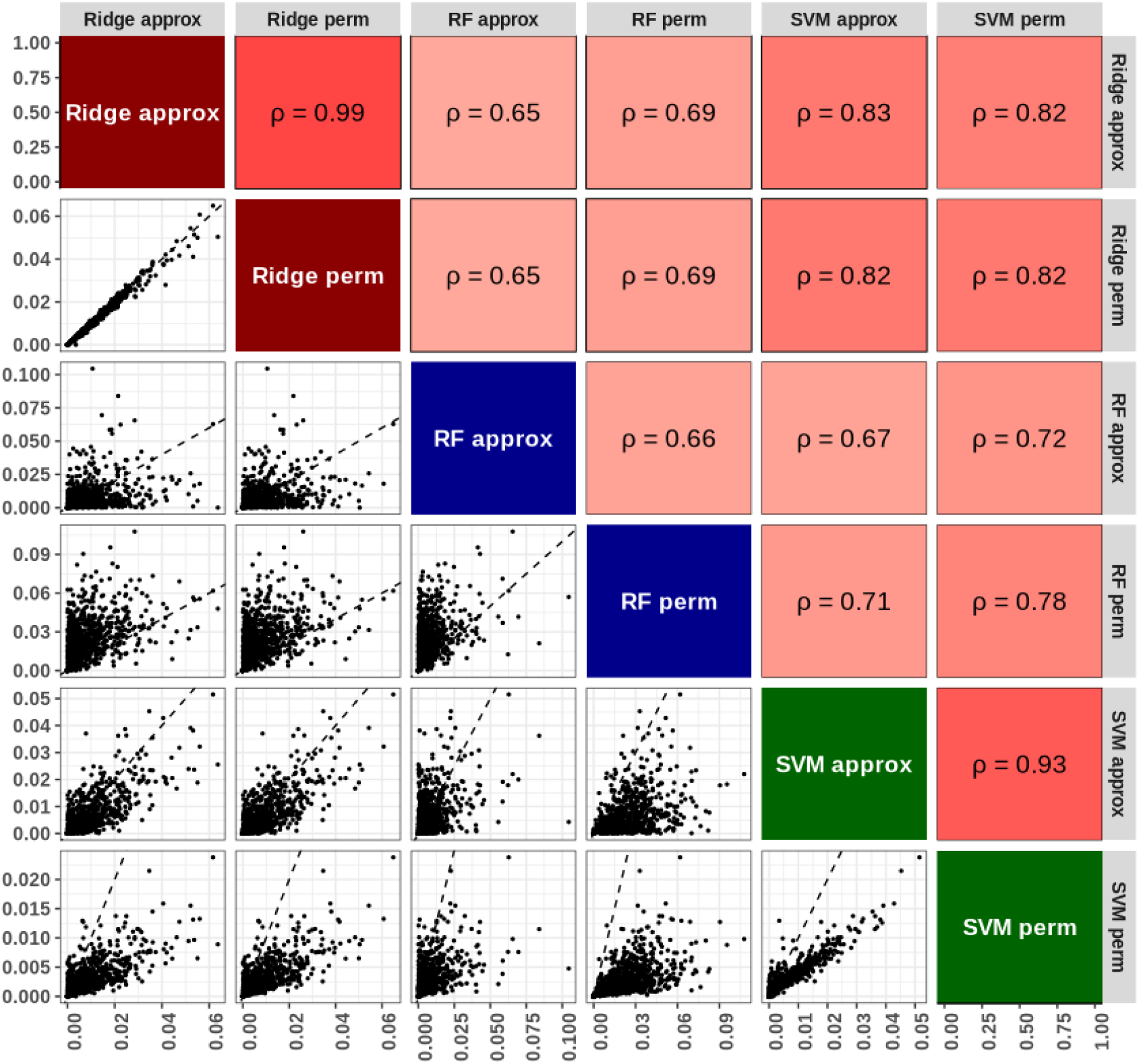
Evaluate linear approximation of permutation feature importance as the co-expression measure (Supplementary Results S1.1). Biplot assessing the concordance between permutation feature importance (PFI) scores derived using the linear approximation described in Supplementary Methods S3.3.2 and those calculated via actual permutation. Lower triangle: scatter plots of PFI scores computed with different regression models (Ridge: ridge regression, RF: random forest, SVM: support vector machine), comparing the linear approximation and actual permutation. Dashed lines represent dentity lines. Upper triangle: correlation coefficients between PFI scores derived from different approaches. The figure demonstrates high correlation between approximated and actual PFI scores for linear models and SVMs, showcasing the accuracy and computational efficiency of the approximation

**Figure S5.**
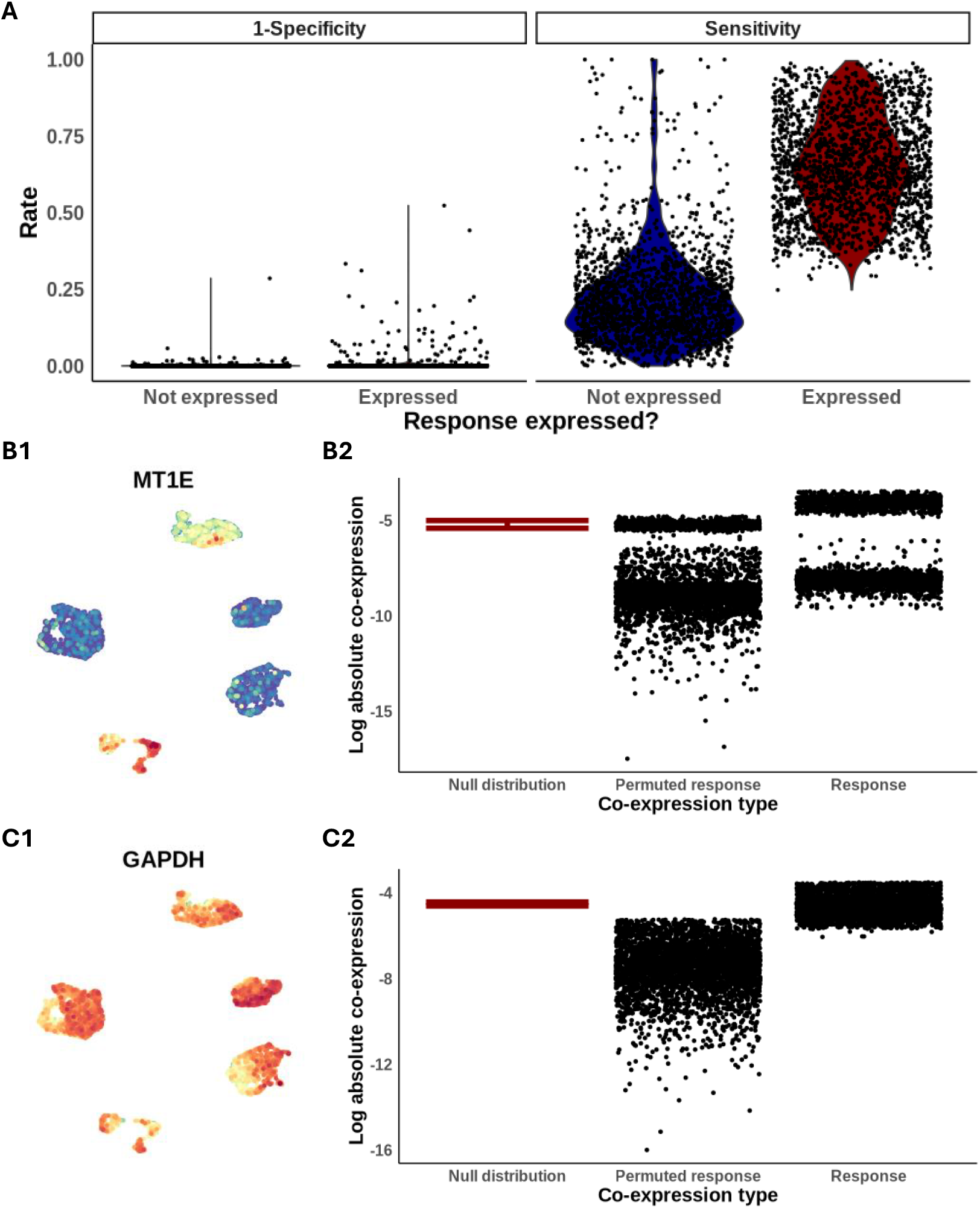
Sensitivity and Specificity Analysis of NNet Pruning Based on Significance Measures (Supplementary Results S1.2). **(A)** A specialized dataset was constructed for this analysis, comprising the 2,000 most variable features (VFs) from the human lung adenocarcinoma cell line dataset of Tian et al. [2019], along with a permuted counterpart for each VF. Using NNet, we evaluated each VF as a response gene and assessed the pruning effectiveness in two aspects: sensitivity (the ability to retain a response gene’s co-expression with itself as significant) and specificity (the ability to filter out the response gene’s co-expression with its permuted counterpart). The analysis was conducted separately on cell populations expressing and not expressing the response gene. Results are visualized as violin plots, with each jittered scatter point representing the outcome for a specific response gene.**(B)** An example showcasing the response gene MT1E, where NNet pruning successfully constructed a null distribution of co-expression that distinguished self-co-expression from co-expression with the permuted counterpart. (B1) The expression of MT1E across the dataset. (B2) The distribution of co-expression values illustrating the separation between significant and insignificant co-expression. **(C)** Similar to (B), but showing an example of a housekeeping gene, GAPDH, as the response gene. In this case, NNet pruning was overly aggressive, classifying a large proportion of self-co-expression as insignificant, potentially due to the uniform expression patterns of housekeeping genes.

## S3 Supplementary Methods

## S3.1 Abbreviations

scRNA-seq: single cell RNA-sequencing
TF: transcriptional factor
PC: principal component
KNN: k-nearest neighbours
PCA: principal component analysis
nPCA: non-negative principal component analysis
UMAP: uniform manifold approximation and projection
PPR: personalised page-rank
LRA: low-rank approximation
NTF: non-negative tensor factorisation
NMF: non-negative matrix factorisation
SVD: singular value decomposition
PKN: prior knowledge network
USP: upperstream siganling pathway
PLS: partial least square regression

## S3.2 An overview

NNet builds a gene co-expression (or regulatory) network for each cell based on single-cell RNA-seq (scRNA-seq) data. The methods consists of two major components:

1. Building cell-specific networks (Section S3.3). This component infers gene co-expression using principal component (PC) regression within each cell’s k-nearest neighbours (KNNs).
2. Downstream analysis. This includes meta-network construction (Section S3.4) and function annotation (Section S3.5) of the inferred co-expression networks using prior knowledge in gene regulation and signalling interactions.

Additionally, Section S3.6 describes a subsampling method to address the memory usage challenges associated with loading and analysing large networks of thousands of cells.

We briefly describe the PC regression and prior network annotation below:

### S3.2.1 Cell-specific KNN-PC regression

A detailed description of the method can be found in Section S3.3. Here we present an abstract of the method in a sequential order. Steps that are performed globally on the full data and locally on each cell’s KNN are noted as *Global* and *Local* respectively.

1. (Global) Principal component analysis (PCA) on the full scRNA-seq data (Section S3.3.1).
2. (Global) Construct a cell k-nearest neighbour graph based on PCA. The graph defines the neighbouring cells of each cell and their similarity in gene expression. The local co-expression network for each cell is then learned within its defined neighbourhood.
3. **For each gene in a response gene sets, repeat step 4 to 6**
4. (Local) Within each cell’s neighbourhood, fit a PC regression model on the expression of the response gene.
5. (Local) On each model fitted, calculate importance of genes in predicting the response based on permutation feature importance. The importance score of a gene serves as a measure of its co-expression with the response at the corresponding cell level (Section S3.3.2).
6. (Global) Apply random walk diffusion to denoise the cell by predictor gene importance score matrix of the response (Section S3.3.4).

On each cell we measure co-expression between multiple response genes and predictor genes, which builds a co-expression network. Each cell has its own network (i.e, cell-specific network), and the collection of networks builds a network tensor (a cell-by-gene-by-gene data entity storing networks, see Section S3.3.3 for details). A optional downstream analysis is applying non-negative PCA (nPCA) on cell space of the network tensor to extract meta-networks that represent principal co-expression patterns (Section S3.4).

### S3.2.2 Prior knowledge annotation of co-expression networks

A detailed description of the method can be found in Section S3.5. We have two major goals in this part of method development. The first is to discovery context-specific activation of gene regulation. Genes that are both known to interact based on prior knowledge and are co-expressed in the study context are more likely to represent on-going gene regulation. The second goal is to identify upstream signalling pathways and receptors that could potentially influence the expression of a target gene, given its co-expression with transcription factors (TF). To achieve these goals, we

1. (Pre-computed) Constructed large prior knowledge networks of gene regulation (TF-targets) and protein-protein interaction integrated from multiple confidential databases [Türei et al., 2021]. The integration approach we took was acquiesced from Browaeys et al. [2020]. The prior knowledge network is included in the R package.
2. (Pre-computed) Build a receptor-TF regulatory potential matrix by running personalised page-rank algorithm (PPR) on the prior knowledge network.
3. (Local) Quantify receptors’ regulatory activity on targets by integrating prior knowledge in receptor-TF regulatory potential with target-TF co-expression. Receptors that strongly correlates with a target through TFs are more likely to be the signalling transductor of target gene expression.

## S3.3 Cell-specific KNN-PC regression

We infer gene co-expression using regression models, which quantify the relationships between genes by estimating the importance of predictor gene expression in predicting response gene expression. Cell-specific co-expression is determined by fitting regression models to each cell’s k-nearest neighbours. A significant technical challenge we face is the sparsity, noise, and immense size of scRNA-seq data, which can lead to inaccurate results and make the analysis non-scalable. To address this challenge, we propose a solution using PC regression.

### S3.3.1 PC regression

Most existing GRN inference methods calculate gene co-expression directly within the gene expression space, involving computations across tens of thousands of gene pairs [Langfelder and Horvath, 2008, Huynh-Thu et al., 2010, Meyer et al., 2007]. Performing this for each cell’s neighbourhood is computationally infeasible.

PC regression is an attractive solution to the computational challenge we face [Jolliffe, 1982]. Instead of fitting responses directly on gene expressions, we fit on the PC of gene expression. PCA not only speeds up model fitting by embedding data onto a much lower dimension, but also internally denoises and imputes scRNA-seq since a low-rank approximation (LRA) of the data can be recovered from PC. We derive gene-level importance of a PC regression model by estimating permutation feature importance (Section S3.3.2). In order to perform PC regression, we first run PCA on the full data including every cells

#### PCA

Let 𝒳 be a *N* by *P* scRNA-seq gene expression matrix that is centered to zero mean and scaled to a unit variance. We extract the first *R* < *min*(*N, P*) singluar vector of 𝒳 to construct a rank-*R* LRA of 𝒳 as

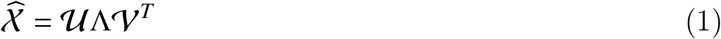

Here, the *N* × *R* matrix 𝒮 = 𝒰Λ is known as the scores, and the *P* × *R* matrix 𝒱 is known as the loading vectors in PCA.

We choose *R* according to the method described in [Linderman et al., 2022], who estimate *R* by evaluating the spacing between singular values *s*_*r*_ = *λ*_*r*+1_ − *λ*_*r*_, where *λ*_*r*_ = Λ_*rr*_. *R* is chosen as the largest *r* ∈ {1, …, 100} such that *s*_*r*_ is smaller than a specific threshold determined by the distribution of {*s*_*r*_ : *r* ∈ {1, …, 100}}. See the original paper for details.

#### Construct KNN-graph

We begin by constructing a KNN graph of cells based on the distances between them in the PCA space. This KNN graph serves multiple purposes: it facilitates the execution of weighted PC regression, enables the calculation of weighted local variance of gene expression (as described in Section S3.3.2), and assists in denoising the co-expression estimation (detailed in Section S3.3.4).

The KNN-graph is constructed according to the uniform manifold approximation and projection algorithm, as outlined by Becht et al. [2019]. Below, we describe the *N* × *N* weighted adjancency matrix 𝒲 that represents the KNN-graph. The detailed reasoning for choosing this KNN-graph is similar to that discussed in [Deng et al., 2022], and we omit it here for brevity.

In the KNN graph, each cell *n* ∈ {1, …, *N*} is connected to its *K*-nearest neighbours, represented as a set KNN(*n*) ⊆ {1, …, *N*}. The (*n, m*) entry of 𝒲, which represents the weight of the edge directed from cell *n* to cell *m* on the KNN graph, is given by:

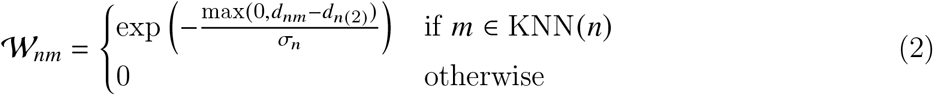

where *d*_*nm*_ = ∥𝒮_*n*._ − 𝒮_*m*._ ∥ is the Euclidean distance in PC scores between cells *n* and *m*. The notation *d*_*n*(*k*)_ denotes the distance from cell *n* to its *k*^*th*^ nearest neighbor. Specifically, *k* = 1 refers to cell *n* itself, while *k* = 2 refers to the nearest neighbor of cell *n*. The scaling factor *σ*_*n*_ is chosen such that each node has a fixed out degree equals to log_2_ (*K*),

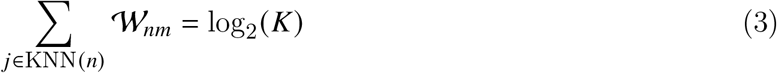

In the following sections, we will use 𝒲_*n*(*k*)_ to denote the edge weight between cell *n* and its *k*^*th*^ nearest neighbour, similar to the notation of *d*_*n*(*k*)_.

#### PC regression

We describe PC regression in cell *n*’s *K*-nearest neighbourhood. To simplify notations, we use bold letter 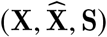 to distinguish local data from that of global ones 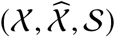 respectively, without mentioning *n*.

Let 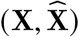 be *K*×*P* gene expression data sampled at cell *n*’s *K*-nearest neighbourhood. The *K*×*R* matrix **S** and the length *K* vector 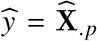 represents the PC scores and the response *p*, respectively, for the cell neighbourhood. 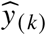 and **S**_(*k*)._ for *k* ∈ {1, …, *K*} represents the response and the PC score of the *k*^*th*^ nearest neighbour of cell *n* respectively.

We weight cells in the neighbourhood of cell *n* according to the KNN-graph we constructed in the privous section. PC regression with data of the *k*^*th*^ neighbour (*y*_(*k*)_, **S**_(*k*)._) weighted by 𝒲_*n*(*k*)_ is then performed,

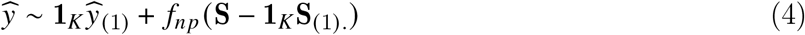

where *f*_*np*_ is a regression model of choice. Since we are only interested in the local partial effect of PCs or genes on the response at cell *n*, we center the regression to cell *n* by adding a intercept term 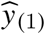 and centering **S** by **S**_(1)._. Our default choice of regression model is ridge regression with a penalty parameter set to 0.5. The solution of this ridge regression is equivalent to the maximum a posteriori solution of a Bayesian regression with standard normal priors on the regression coefficients, which is a prior choice suggested by [Bürkner, 2017].

### S3.3.2 Permutation feature importance as co-expression measure

We first introduce the feature importance score for predictor gene *q* on response gene *p*, at cell *n*, denoted as 𝒜_*npq*_. This score is based on an approximation of permutation feature importance from eq. (12) and adjusted for bias in PC regression. By construction, 𝒜_*npq*_ is the square of the effect measure ℬ_*npq*_, which can be intepreted as the change in the response gene expression resulting from a standard unit change in the predictor gene expression.

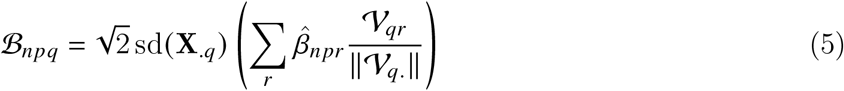

and

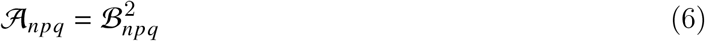

Here, sd(**X**_.*q*_) is the local weighted standard deviation of the scaled predictor gene *q*, weighted by 𝒲_*n*(*k*)_. The term 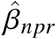 is the partial derivative of the fitted regression function 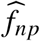 with respect to the *r*^*th*^ principal component at cell *n*. In practice, our method returns ℬ_*npq*_ instead of 𝒜_*npq*_ because the former retains the sign of the effect, providing more informative results. 𝒜_*npq*_ can be easily derived from ℬ_*npq*_ when needed

The following sections detail the derivation and rationale behind our feature importance score.

#### Permutation feature importance

We propose to measure co-expression between a response and predictor genes by the importance of predictor genes in fitting regression model. Since we do not want to impose parametric assumption on *f*, we consider using permutation feature importance, a commonly used predictor importance score in non-parametric regression. In our PC regression context, predictor *q*’s importance is calculated by empirically estimating the following expectation

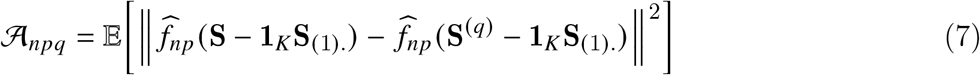

where 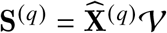 is the PC score projection with the *q*^*th*^ gene of 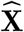 being resampled locally in the cell neighbourhood. 𝒜_*npq*_ represents the mean squared difference between the original PC regression prediction, and the prediction made subject to resampling of gene *q*. A larger value of 𝒜_*npq*_ indicates that the gene is more influential in the regression model. Typically, 𝒜_*npq*_ is estimated by permuting the gene data and calculating the squared differences for multiple times, then averaging these differences over all permutations.

#### A simplified approach for estimation

Permutation approach for estimating 𝒜_*npq*_ is computationally expensive, especially when we need to fit regression and permute for each gene in each cell’s neighbourhood. To address this, we propose a simplified approach for estimating 𝒜_*npq*_. We numerically validated our proposed estimator on different PC regression models.

Let **X** be samples from a random vector *X* of length *P*, representing the distribution of gene expression in the cell neighbourhood. Define *X* ^(*q*)^ as a variant of *X*, with 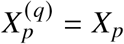 for all *p* ≠ *q*. 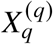 is sampled from the same distribution as *X*_*q*_, but independent from *X*_*q*_. eq. (7) in this population setting can be simplified as

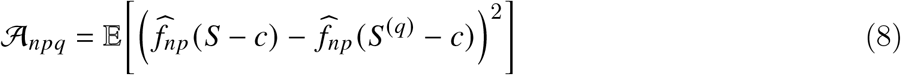

where *S* = *X*𝒱 and *S*^(*q*)^ = *X*^(*q*)^𝒱 are length *R* random vectors representing local PC score distributions, *c* is a constant centering vector. Using a first-order Taylor expansion around 0, we obtain

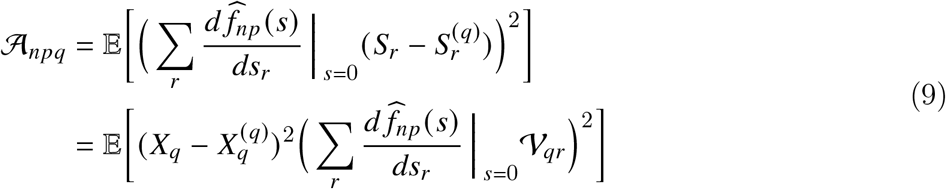

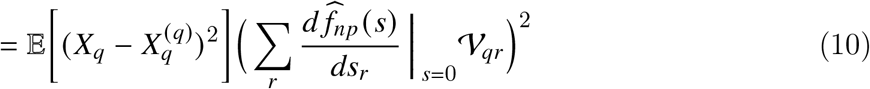

When centreing regression around cell *n*, 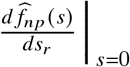 represents the partial derivative of 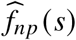 evaluated at cell *n* since *s* = *S* − *c* = 0. We simplify the notation of these partial derivative as 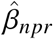. In the case of a simple linear regression model, the derivative 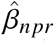 is the regression coefficient fitted on the *r*^*th*^ component. In the cases of non-linear regressions, 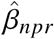 is estimated using symmetric derivative of moving average smoothed 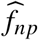 [Aull, 1967, Jacoby, 2000].

For calculating the first expectation term of eq. (10), we leverage the iid assumption of *X*_*q*_ and 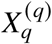 to derive

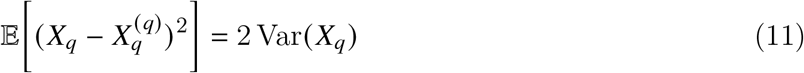

This expression represents twice the local variance of gene *q*, which can be computed directly without the need for permutations. Thus, the expression for 𝒜_*npq*_ becomes:

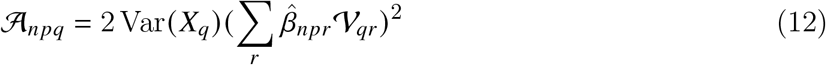

𝒜_*npq*_ can now be easily interpreted as we introduced at the beginning of Section S3.3.2, hence provides a straightforward measure of feature importance.

To evaluate the accuracy of the approximation in eq. (12) for 𝒜_*npq*_, we compared it against values of 𝒜_*npq*_ obtained via actual permutation, as defined in eq. (7) (Supplementary Results S1.1). For linear models, including linear support vector machines and ridge regression, we achieved nearly perfect approximations. In the case of non-linear models like random forests, which have been successful in inferring gene co-expression [Huynh-Thu et al., 2010], we observed correlations exceeding 0.5 between results of approximation and permutation.

#### Adjust bias in PC regression

Note that 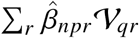 in eq. (12) represents the dot product between 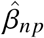. and 𝒱_*q*._. This dot product is proportional to ∥𝒱_*q*._∥, which is the magnitude of the loadings for gene. Consequently, genes that are more influential in the PCA tend to receive higher importance scores. However, we contend that ∥𝒱_*q*._∥ should not significantly influence the importance score. What truly matters is whether 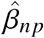. and 𝒱_*q*._ are aligned in the same direction.

To address this bias, we propose scaling 𝒱_*q*._ to a unit length. This adjustment leads to our final expression for the sample-level importance score, as proposed in eq. (5).

### S3.3.3 Matrix representation of networks and network tensor

When performing regression on *P*′ ≤ *P* response variables, we obtain a *P*′ ×*P* importance score matrix 𝒜_*n*.._, for each cell *n* ∈ 1, …, *N*. This matrix 𝒜_*n*.._ serves as a representation of the co-expression network for cell, where each element reflects the strength of the connection (i.e., co-expression calculated as described in eq. (5)) between pairs of nodes (i.e, genes).

Building on this concept, the *N* ×*P*′ ×*P* data entity 𝒜, known as a network tensor, is a collection of the co-expression networks across all cells. Similarly, ℬ, which represents a stack of effect matrices, is referred to as an effect tensor.

### S3.3.4 Graph denoise by random walk diffusion

Regression can be unstable and noisy when there is a small sample size in each cell’s neighbourhood. Suppose we construct an *N* × *P*′ × *P* effect tensor ℬ. To denoise the effect estimation, we smooth ℬ over the KNN-graph of cells that is described in S3.3. We first symmetrise 𝒲 as done by [Becht et al., 2019]

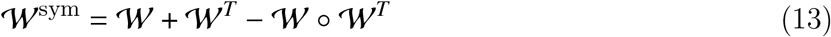

where ° denotes the Hadamard product. Next, we construct a random walk operator:

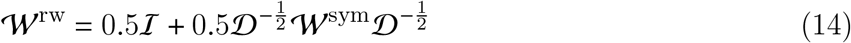

where ℐ is an identity matrix, 𝒟 is a diagonal matrix with 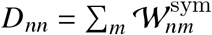. The largest eigenvalue of 𝒲^rw^ is equal to 1, and the remaining eigenvalues are bounded by 0 and 1. This construction ensures that the diffusion process does not drift the scale of ℬ. The effect matrix of cell *n*, ℬ_*n*.._, is denoised by applying the random walk operator on ℬ:

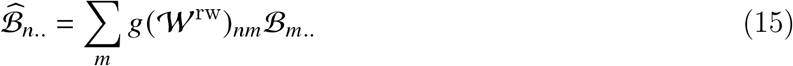

where *g* is a function on the eigenvalues of 𝒲^rw^, given by:

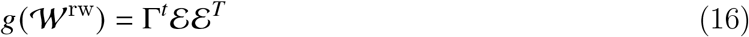

Here, Γ are the eigenvalues and ℰ are the corresponding eigenvectors of 𝒲^rw^. *t* is a hyper-parameter that controls the number of steps of the random walk. If Γ and ℰ represents the complete eigensystem of 𝒲^rw^, we have *g*(𝒲^rw^) equals to the *t*^*th*^ power of 𝒲^rw^. In practice, we only construct an eigensystem with a rank much smaller than *N*, allowing the diffusion to be more efficiently computed in eq. (15) via matrix multiplication. Moreover, the use of a rank-reduced *g*(𝒲^rw^) can be interpreted as applying low-pass filtering on ℬ [Smola and Kondor, 2003], resulting in a smoother 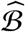 over the graph.

The smoothed importance score 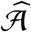 can be calculated as an element-wise square of 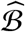.

### S3.3.5 Network pruning

The denoised effect tensor 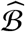 is fully dense and contains a substantial amounts of insignificant effects that do not represent true co-expression. To prevent these insignificant effects from confounding the downstream analysis, we prune 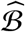 independently for each response gene by calculating the statistical significance of effects and potentially shrink non-significant effects to zero. We describe in here the pruning of effects of *P* predictor genes on a specific response gene *p*, which is represented by the effect matrix 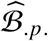 of size *N* ×*P*.

The pruning strategy is based on the premise that response genes can act as their own predictors in the PC regression model. Given that a gene must be a causal regulator of itself, the effect of the response on itself, denoted by 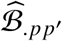, where *p*′ is the index of the response as a predictor, provides a benchmark. This benchmark can identify effects that might indicate functional co-regulation with the response.

Specifically, let ℬ_.*p*._ represent the noisy co-expression with the response *p* and 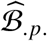 the estimated signals derived from ℬ_.*p*._. To establish a null distribution of signals, we conduct random diffusion by permuting the entries (cells) in ℬ_.*pp*′_ to create a vector of destructed effects, denoted by *ω*_*p*_ Picart-Armada et al. [2021]. Subsequently, we apply the diffusion operator *g*(𝒲^rw^) as defined in eq. (15) to denoise *ϵ*_*p*_:

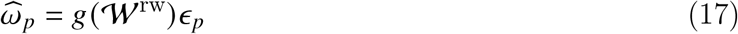

The resulting vector 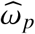 represents the signals from random data, which we consider as indicative of insignificance. We repeat the permutation and calculate the mean 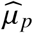 and the standard deviation 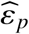 of log 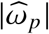 over permutation. The significance of 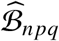, representing the effect of gene *q* in predicting the response in cell *n*, is given by

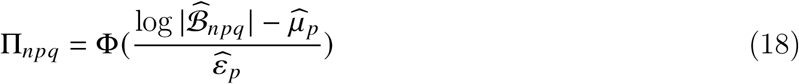

Here, the left-hand side of the equation represents the lower tail probability of log 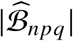 in a normal distribution 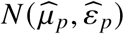. In this context, Φ denotes the cumulative distribution function of the standard normal. We classify 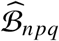 as significant if it deviates from the distribution of 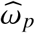, thereby receiving a larger value of Π_*npq*_ closer to 1.

A *N* × *P*′ × *P* tensor of significant scores Π is constructed by repeating the above procedure for each *p* ∈ {1, …, *P*′}.

## S3.4 Downstream analysis: embed network ensemble by non-negative PCA

After obtaining the denoised importance score tensor 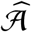, which represents cell-specific networks, an optional downstream analysis involves discovering the most informative network patterns in the data. This can be achieved by embedding the cell dimension using methods such as non-negative tensor factorisation (NTF).

However, given the potentially large size of 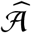, NTF may not be computationally feasible in terms of speed and memory usage. To address this, we simplify the procedure by applying non-negative matrix factorisation (NMF) to a matrix representation of 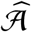. This is accomplished by vectorizing the adjacency matrix of each cell, 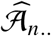, into a one-dimensional vector of edge weights. The resulting tall-and-skinny matrix should have *P*′*P* >> *N* rows corresponding to edges between *P*′ responses, and *P* predictors and *N* columns corresponding to cells.

We adapted the approach taken by Benson et al. [2014] to efficiently solve the NMF problem for tall-and-skinny matrices. At the core of this approach is the reduction of the row dimension of the matrix using singular value decomposition (SVD). Instead of applying NMF to the entire matrix, it is applied to the singular vectors obtained from SVD, which significantly reduces the computational complexity. We found that the approach is also valid when non-negative PCA (nPCA) is the NMF method.

We consider solving nPCA with the objective of maximising variances of PC scores subject to the non-negativity constrain on loadings.

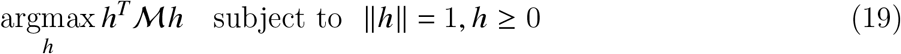

where the covariance matrix between cells ℳ can be calculated by

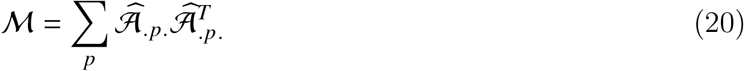

We observe that by diagnosing ℳ = ℒΣℒ, we have

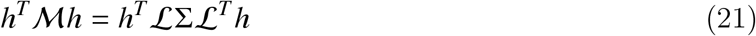

Combining with equation (21), a straightforward derivation shows that the objective in equation (19) is equivalent to solving a restricted singular value decomposition (SVD) problem on 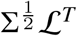 [Shen and Huang, 2008], which has much lower dimension compared to the tall-and-skinny matrix previously described. Additionally, it eliminates the need to construct the tall-and-skinny matrix, thereby saving on extra memory usage. This SVD problem can be efficiently solved using iterative regression and deflation techniques, as detailed in Mackey [2008] and Sigg and Buhmann [2008]. Without details in derivation, we describe the algorithm in below.

### Algorithm 1 Non-negative principal component analysis (nPCA) on long and tall matrices

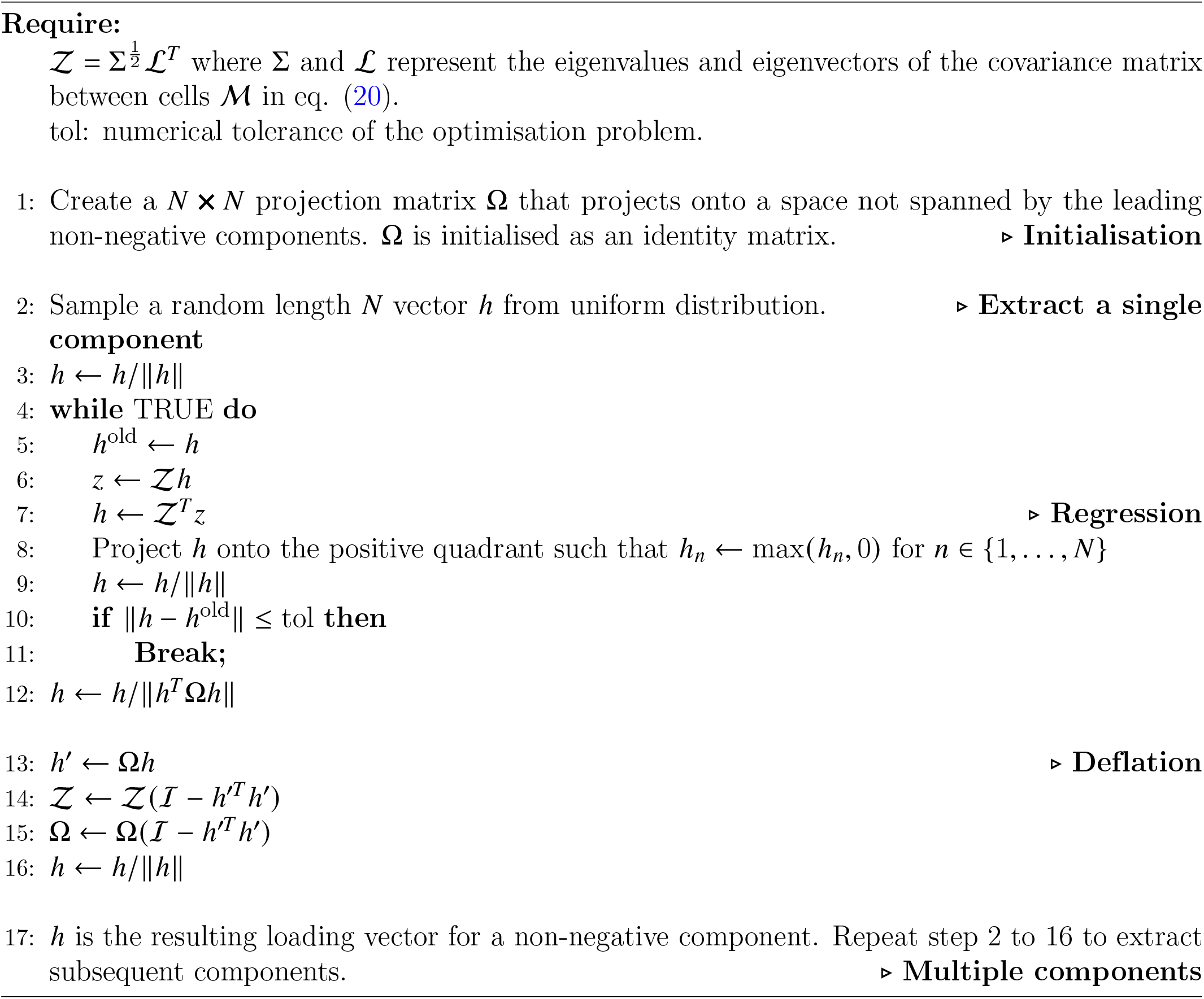

To further improve computational efficiency in practice, we truncate 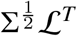 by retaining only the leading eigenvalues in 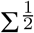. nPCA is performed on the truncated matrix.

## S3.5 Downstream analysis: prior knowledge annotation of co-expression networks

Now we have a tensor 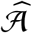 (see Section S3.3) that represents cell-specific co-expression networks. These networks are termed ‘co-expression’ because they indicate observed associations between gene expressions; however, they do not necessarily represent direct gene regulation. For instance, two genes may display highly correlated expression patterns without directly interacting with each other. Such co-expression can be confounded by common upstream regulators. To improve the interpretability of our co-expression networks, our next task is to annotate them with prior knowledge regarding the direction of gene regulation. Genes that are correlated in expression and supported by prior evidence provide a more reliable representation of biologically relevant, context-specific regulatory mechanisms.

In this section, we first describe how we construct a confidential prior knowledge network (PKN) by integrating multiple gene regulation and signalling interaction databases using the R package `OmniPath` [Türei et al., 2021] (Section S3.5.1). We then explain how we incorporate this prior knowledge into our co-expression network (Section S3.5.2).

### S3.5.1 Construct prior knowledge network with OmniPath

OmniPath is an R package that serves as an interface to an online database of prior knowledge in intra- and inter-cellular signalling interactions. It also provides pipelines for integrating user-chosen databases and quality controls to build custom PKN. We build our network with OmniPath version 3.8.2, and adapated the built-in NicheNet integration pipeline originally proposed by Browaeys et al. [2020].

#### Network resources

We run `OmniPath::nichenet_networks` to extract public inter- and intra-cellular signalling networks, and gene regulatory networks. We only consider direct interaction networks curated by Omnipath. For inter-cellular signalling networks, we only consider ligands and secreted enzymes as parental nodes, with receptors and transporters as child nodes. For the gene regulatory network, we exclusively use the CollecTRI database, which is an integrated database of high-quality evidence for TF and target interactions [Müller-Dott et al., 2023].

#### NicheNet Prior Knowledge Integration with Omnipath

NicheNet is a popular method that models ligand and target gene interactions using prior-knowledge networks Browaeys et al. [2020]. In the paper, the authors describe a method to integrate networks from different sources using automatic parameter optimization, with a objective of building an integrated network that best aligns with ligand perturbation experiments. We implement NicheNet integration with Omnipath, following the tutorial at https://r.omnipathdb.org/articles/nichenet.html (Last retrieved: July 30th, 2024).

The integration process yields a weighted gene regulatory network, a weighted signalling interaction network that encompasses both inter- and intra-cellular signalling pathways, and a matrix summarising ligands’ regulatory potential on the other genes. These integrated data sets are provided by our package and are readily available for user application.

#### Receptor-target regulatory potential

After the NicheNet integration, a ligand regulaory potential matrix is constructed by applying the personalized PageRank (PPR) algorithm to the signalling interaction PKN [Page, 1999]. Specifically, the regulatory potential computed by PPR represents the probability that a signal, originating from a ligand, is transmitted to the target in the steady state of a random walk on the PKN.

In a similar manner, we construct a receptor regulatory potential matrix. We implement PPR using `igraph::page_rank` function from the R package `igraph` (version 2.0.1.1) [Csardi and Nepusz, 2006]. Other than setting receptors as the source nodes, the parameters for running PPR are the same as those optimized during the integration process for the ligand regulaory potential matrix.

### S3.5.2 Prior knowledge annotation

Here, we focus on the annotation for co-expression networks between TFs and potential target genes. In this context, one set (either TFs or targets) as responses and the other as predictors. The underlying assumption is that gene regulation requires co-expression between TFs and targets. In contrast, receptors do not need to be co-expressed with their targets to transmit signals; they only need to be expressed.

#### Annotate Co-Expression Networks

We categorise edges (i.e., co-expression) in 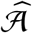 into four confidence levels. These levels will be visually distinguished in the network visualisation function provided by our package. The four confidence levels are described in an increasing order below:

- Non-significantly co-expressed: The significance of an edge is calculated according to S3.3.2. These edges are not be shown in the visualisation.
- Significantly co-expressed, but lacking evidence of physical interaction: These edges are classified as significant, but there is no existing evidence in the prior gene regulatory network to support that the co-expression is regulatory. These edges are represented as dashed lines in the co-expression network.
- Significantly co-expressed and interacting within a few steps of signal transduction: These edges are classified as significant, and the TF signal can reach the target within a few steps (as user defined) in the prior gene regulatory network. These edges are represented as indirected solid lines in the co-expression network.
- Significantly co-expressed and directly interacts: These edges are classified as significant, and the corresponding TF directly interact with the target (in one step) in the prior gene regulatory network. These edges are represented as directed solid arrows in the co-expression network. Activation and suppression regulation are distinguished by the arrow heads.

#### Infer potential up-stream signalling pathways of target genes

We describe a scenario where we run KNN-PC regression using target genes as responses, and aim to infer up-stream signalling pathways or receptors potentially affecting target expression in a context-specific manner. We begin by measuring receptor activity on the targets by combining co-expression networks with a prior model of receptor potential, created using the PPR algorithm (see Section S3.5.1). A receptor *i*’s activity in regulating a single target *p* in cell *n* is quantified by

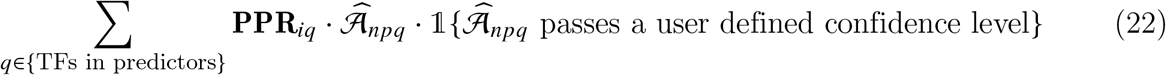

where **PPR**_*iq*_ represents the receptor’s regulatory potential on TF *q*. Summand in the equation reflects the receptor’s inverse regulatory distance to the target, mediated by TFs, and is further pruned by the confidence of 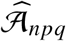.

For multiple targets, we can perform integrative analysis between the receptor potential matrix **PPR** and the cell *n*’s co-expression network 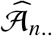, using tools such as those in the mixOmics package [Rohart et al., 2017]. We describe a partial least square regression (PLS) [Wold et al., 2001] approach that is a multivariate analogue to eq. (22). We first extract relevant data seubsets **PPR**^TF^ and 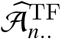 that have matching columns that represents known TFs in the predictors. The PLS algorithm finds linear embeddings for receptors and targets that maximize the covariance of TFs in this embedded space:

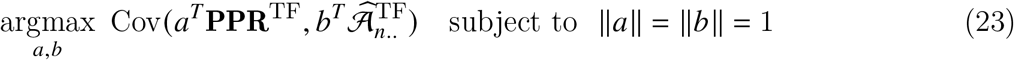

Intuitively, *a* and *b* represent receptor and target modules that are most strongly connected through TFs. Receptors that are highly weighted by *a* are more likely to actively regulate the targets.

Finally, we infer the potential signalling pathway between a likely active receptor *i* and a target *p*. To do this, we first identify the TF *q* through which the distance between the receptor and the target is the shortest. Specifically, this TF is found by

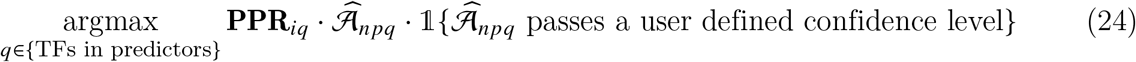

We simply consider the potential signalling pathway as the shortest path between TF *q* and the receptor, which is searched on the prior signalling network by the R igraph function `igraph::shortest_paths`.

## S3.6 Analysis on subsampled data

Up to now, we have presented a computationally efficient algorithm for inferring cell-specific co-expression networks. However, we have not yet addressed the issue of memory usage. Storing dense networks for tens of thousands of cells is impractical on a personal computer and can be costly on a server.

We propose to address the memory usage problem by implementing parts of our pipeline on only a subset of cells that are representative of the full dataset. The following steps are performed on the entire dataset:

- PCA, LRA and KNN-graph construction.

For the representative subset of cells, we conduct:

- PC regression, importance score calculation, graph denoise and nPCA to construct meta-network.

### Select representative cells

To select representative cells, we perform k-means clustering on the PC scores *S* with *N*′ centers. Within each cluster, we choose the cells most similar to the cluster center as representative cells for that cluster. Other sophisticated method, such as MetaCell [Baran et al., 2019], for selecting representative cells can be integrated seamlessly into our method. Next we describe the analysis on representative cells in more details.

### PC regression and importance score calculation

For a representative cell, its neighbours, the weighting of neighbours, and the regression data are consistent with the full analysis. Thus, the procedure is equivalent to applying the methods described in Sections S3.3.1 and S3.3.2 for each cell *n* ∈ 1, …, *N*, while skipping those non-representative cells. Ultimately, this process yields a smaller effect tensor ℬ′ of size *N*′ ×*P*′ ×*P* that is a subset of ℬ obatined from the full analysis.

### Graph denoise

We denoise ℬ′ and, if needed, impute the effects of non-representative cells using network propagation. First, we construct an *N* ×*N*′ weight matrix 𝒲′, which extracts columns of *g*(𝒲^rw^) defined in eq. (16), corresponding to the representative cells in ℬ′. We then further normalize 𝒲′ by rows so that each row sums to one. The denoised (or impute) effect of cell *n* 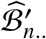 for *n* ∈ {1, …, *N*} is computed by

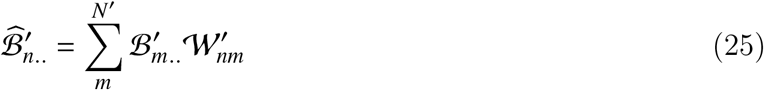

### Network pruning

The null distribution of effect size for a response *p* is learned analogously as in S3.3.5 using 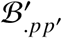, which is the response’s effect on itself across representative cells. Cells, including non-representing ones, are pruned by comparing 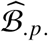 with the null distribution, as done in eq. (18).

### nPCA

nPCA is conducted on the denoised importance score tensor of representative cells.

## References

Vân Anh Huynh-Thu, Alexandre Irrthum, Louis Wehenkel, and Pierre Geurts. Inferring regulatory networks from expression data using tree-based methods. PloS one, 5(9):e12776, 2010.

Peter Langfelder and Steve Horvath. Wgcna: an r package for weighted correlation network analysis. BMC bioinformatics, 9(1):1–13, 2008.

Pau Badia-i Mompel, Lorna Wessels, Sophia Müller-Dott, Rémi Trimbour, Ricardo O Ramirez Flores, Ricard Argelaguet, and Julio Saez-Rodriguez. Gene regulatory network inference in the era of single-cell multi-omics. Nature Reviews Genetics, 24(11):739–754, 2023.

Thalia E Chan, Michael PH Stumpf, and Ann C Babtie. Gene regulatory network inference from single-cell data using multivariate information measures. Cell systems, 5(3):251–267, 2017.

Shilu Zhang, Saptarshi Pyne, Stefan Pietrzak, Spencer Halberg, Sunnie Grace McCalla, Alireza Fotuhi Siahpirani, Rupa Sridharan, and Sushmita Roy. Inference of cell type-specific gene regulatory networks on cell lineages from single cell omic datasets. Nature Communications, 14(1):3064, 2023.

Samuel Morabito, Fairlie Reese, Negin Rahimzadeh, Emily Miyoshi, and Vivek Swarup. hdwgcna identifies co-expression networks in high-dimensional transcriptomics data. Cell reports methods, 3(6), 2023.

Kenji Kamimoto, Blerta Stringa, Christy M Hoffmann, Kunal Jindal, Lilianna Solnica-Krezel, and Samantha A Morris. Dissecting cell identity via network inference and in silico gene perturbation. Nature, 614(7949):742–751, 2023.

Ziqi Zhang, Jongseok Han, L. Song, and Xiuwei Zhang. Cespgrn: Inferring cell-specific gene regulatory networks from single cell multi-omics and spatial data. bioRxiv, pages 2022–03, 2022.

Hao Dai, Lin Li, Tao Zeng, and Luonan Chen. Cell-specific network constructed by single-cell rna sequencing data. Nucleic acids research, 47(11):e62–e62, 2019.

Xuran Wang, David Choi, and Kathryn Roeder. Constructing local cell-specific networks from single-cell data. Proceedings of the National Academy of Sciences, 118(51):e2113178118, 2021.

Stephen Y Zhang and Michael PH Stumpf. Learning cell-specific networks from dynamics and geometry of single cells. bioRxiv, pages 2023–01, 2023.

Lingfei Wang, Nikolaos Trasanidis, Ting Wu, Guanlan Dong, Michael Hu, Daniel E Bauer, and Luca Pinello. Dictys: dynamic gene regulatory network dissects developmental continuum with single-cell multiomics. Nature Methods, 20(9):1368–1378, 2023.

Aditya Pratapa, Amogh P Jalihal, Jeffrey N Law, Aditya Bharadwaj, and TM Murali. Benchmarking algorithms for gene regulatory network inference from single-cell transcriptomic data. Nature methods, 17(2):147–154, 2020.

Hung Nguyen, Duc Tran, Bang Tran, Bahadir Pehlivan, and Tin Nguyen. A comprehensive survey of regulatory network inference methods using single cell rna sequencing data. Briefings in bioinformatics, 22(3):bbaa190, 2021.

Pau Badia-i Mompel, Jesús Vélez Santiago, Jana Braunger, Celina Geiss, Daniel Dimitrov, Sophia Müller-Dott, Petr Taus, Aurelien Dugourd, Christian H Holland, Ricardo O Ramirez Flores, et al. decoupler: ensemble of computational methods to infer biological activities from omics data. Bioinformatics Advances, 2(1):vbac016, 2022.

Atray Dixit, Oren Parnas, Biyu Li, Jenny Chen, Charles P Fulco, Livnat Jerby-Arnon, Nemanja D Marjanovic, Danielle Dionne, Tyler Burks, Raktima Raychowdhury, et al. Perturb-seq: dissecting molecular circuits with scalable single-cell rna profiling of pooled genetic screens. cell, 167(7): 1853–1866, 2016.

Sara Aibar, Carmen Bravo González-Blas, Thomas Moerman, Vân Anh Huynh-Thu, Hana Imrichova, Gert Hulselmans, Florian Rambow, Jean-Christophe Marine, Pierre Geurts, Jan Aerts, et al. Scenic: single-cell regulatory network inference and clustering. Nature methods, 14(11):1083–1086, 2017.

Sophia Müller-Dott, Eirini Tsirvouli, Miguel Vazquez, Ricardo O Ramirez Flores, Pau Badia-i Mompel, Robin Fallegger, Dénes Türei, Astrid Lægreid, and Julio Saez-Rodriguez. Expanding the coverage of regulons from high-confidence prior knowledge for accurate estimation of transcription factor activities. Nucleic acids research, 51(20):10934–10949, 2023.

Susanne J Szabo, Sean T Kim, Gina L Costa, Xiankui Zhang, C Garrison Fathman, and Laurie H Glimcher. A novel transcription factor, t-bet, directs th1 lineage commitment. Cell, 100(6):655–669, 2000.

James J Knox, Arpita Myles, and Michael P Cancro. T-bet+ memory b cells: generation, function, and fate. Immunological reviews, 288(1):149–160, 2019.

Danilo Pellin, Mariana Loperfido, Cristina Baricordi, Samuel L Wolock, Annita Montepeloso, Olga K Weinberg, Alessandra Biffi, Allon M Klein, and Luca Biasco. A comprehensive single cell transcriptional landscape of human hematopoietic progenitors. Nature communications, 10 (1):2395, 2019.

Senthil Raja Jayapal, Kian Leong Lee, Peng Ji, Philipp Kaldis, Bing Lim, and Harvey F Lodish. Down-regulation of myc is essential for terminal erythroid maturation. Journal of Biological Chemistry, 285(51):40252–40265, 2010.

M Dolores Delgado and Javier León. Myc roles in hematopoiesis and leukemia. Genes & cancer, 1 (6):605–616, 2010.

Pan Wang, Zi Wang, and Jing Liu. Role of hdacs in normal and malignant hematopoiesis. Molecular cancer, 19:1–21, 2020.

Michael R Tallack, Tom Whitington, Wai Shan Yuen, Elanor N Wainwright, Janelle R Keys, Brooke B Gardiner, Ehsan Nourbakhsh, Nicole Cloonan, Sean M Grimmond, Timothy L Bailey, et al. A global role for klf1 in erythropoiesis revealed by chip-seq in primary erythroid cells. Genome research, 20(8):1052–1063, 2010.

Mira T Kassouf, Jim R Hughes, Stephen Taylor, Simon J McGowan, Shamit Soneji, Angela L Green, Paresh Vyas, and Catherine Porcher. Genome-wide identification of tal1’s functional targets: insights into its mechanisms of action in primary erythroid cells. Genome research, 20 (8):1064–1083, 2010.

Emery H Bresnick, Kyle J Hewitt, Charu Mehta, Sunduz Keles, Robert F Paulson, and Kirby D Johnson. Mechanisms of erythrocyte development and regeneration: implications for regenerative medicine and beyond. Development, 145(1):dev151423, 2018.

Mikiko Suzuki, Maki Kobayashi-Osaki, Shuichi Tsutsumi, Xiaoqing Pan, Shin’ya Ohmori, Jun Takai, Takashi Moriguchi, Osamu Ohneda, Kinuko Ohneda, Ritsuko Shimizu, et al. Gata factor switching from gata 2 to gata 1 contributes to erythroid differentiation. Genes to Cells, 18(11):921–933, 2013.

Laleh Haghverdi, Maren Büttner, F Alexander Wolf, Florian Buettner, and Fabian J Theis. Diffusion pseudotime robustly reconstructs lineage branching. Nature methods, 13(10):845–848, 2016.

Kai Hildner, Brian T Edelson, Whitney E Purtha, Mark Diamond, Hirokazu Matsushita, Masako Kohyama, Boris Calderon, Barbara U Schraml, Emil R Unanue, Michael S Diamond, et al. Batf3 deficiency reveals a critical role for cd8??+ dendritic cells in cytotoxic t cell immunity. Science, 322(5904):1097–1100, 2008.

Ansuman T Satpathy, Xiaodi Wu, Jörn C Albring, and Kenneth M Murphy. Re (de) fining the dendritic cell lineage. Nature immunology, 13(12):1145–1154, 2012.

Jan P Böttcher, Eduardo Bonavita, Probir Chakravarty, Hanna Blees, Mar Cabeza-Cabrerizo, Stefano Sammicheli, Neil C Rogers, Erik Sahai, Santiago Zelenay, and Caetano Reis e Sousa. Nk cells stimulate recruitment of cdc1 into the tumor microenvironment promoting cancer immune control. Cell, 172(5):1022–1037, 2018.

Fábio F Rosa, Cristiana F Pires, Ilia Kurochkin, Evelyn Halitzki, Tasnim Zahan, Nejc Arh, Olga Zimmermannová, Alexandra G Ferreira, Hongzhe Li, Stefan Karlsson, et al. Single-cell transcriptional profiling informs efficient reprogramming of human somatic cells to cross-presenting dendritic cells. Science immunology, 7(69):eabg5539, 2022.

Zahra Elahi, Paul W Angel, Suzanne K Butcher, Nadia Rajab, Jarny Choi, Yidi Deng, Justine D Mintern, Kristen Radford, and Christine A Wells. The human dendritic cell atlas: an integrated transcriptional tool to study human dendritic cell biology. The Journal of Immunology, 209(12):2352–2361, 2022.

Theo W Combes, Federica Orsenigo, Alexander Stewart, AS Jeewaka R Mendis, Deborah Dunn-Walters, Siamon Gordon, and Fernando O Martinez. Csf1r defines the mononuclear phagocyte system lineage in human blood in health and covid-19. Immunotherapy Advances, 1(1):ltab003, 2021.

Daisuke Kurotaki, Michio Yamamoto, Akira Nishiyama, Kazuhiro Uno, Tatsuma Ban, Motohide Ichino, Haruka Sasaki, Satoko Matsunaga, Masahiro Yoshinari, Akihide Ryo, et al. Irf8 inhibits c/ebp?? activity to restrain mononuclear phagocyte progenitors from differentiating into neutrophils. Nature communications, 5(1):4978, 2014.

Roxane Tussiwand and Emmanuel L Gautier. Transcriptional regulation of mononuclear phagocyte development. Frontiers in immunology, 6:533, 2015.

Jacob T Jackson, Yifang Hu, Ruijie Liu, Frederick Masson, Angela d’Amico, Sebastian Carotta, Annie Xin, Mary J Camilleri, Adele M Mount, Axel Kallies, et al. Id2 expression delineates differential checkpoints in the genetic program of cd8??+ and cd103+ dendritic cell lineages. The EMBO journal, 30(13):2690–2704, 2011.

Georgia Anastasiou, Argyri Gialeraki, Efrossyni Merkouri, Marianna Politou, and Anthi Travlou. Thrombomodulin as a regulator of the anticoagulant pathway: implication in the development of thrombosis. Blood Coagulation & Fibrinolysis, 23(1):1–10, 2012.

Matthew Collin and Venetia Bigley. Human dendritic cell subsets: an update. Immunology, 154 (1):3–20, 2018.

Joseph M Chan, Alvaro Quintanal-Villalonga, Vianne Ran Gao, Yubin Xie, Viola Allaj, Ojasvi Chaudhary, Ignas Masilionis, Jacklynn Egger, Andrew Chow, Thomas Walle, et al. Signatures of plasticity, metastasis, and immunosuppression in an atlas of human small cell lung cancer. Cancer cell, 39(11):1479–1496, 2021.

Hui Yu, Liangbin Lin, Zhiqiang Zhang, Huiyuan Zhang, and Hongbo Hu. Targeting nf-??b pathway for the therapy of diseases: mechanism and clinical study. Signal transduction and targeted therapy, 5(1):209, 2020.

Thorsten Hagemann, Subhra K Biswas, Toby Lawrence, Antonio Sica, and Claire E Lewis. Regulation of macrophage function in tumors: the multifaceted role of nf-??b. Blood, The Journal of the American Society of Hematology, 113(14):3139–3146, 2009.

Jingjing Qi, Hongxiang Sun, Yao Zhang, Zhengting Wang, Zhenzhen Xun, Ziyi Li, Xinyu Ding, Rujuan Bao, Liwen Hong, Wenqing Jia, et al. Single-cell and spatial analysis reveal interaction of fap+ fibroblasts and spp1+ macrophages in colorectal cancer. Nature communications, 13(1):1742, 2022.

Ruben Bill, Pratyaksha Wirapati, Marius Messemaker, Whijae Roh, Beatrice Zitti, Florent Duval, Máté Kiss Jong Chul Park, Talia M Saal, Jan Hoelzl, et al. Cxcl9: Spp1 macrophage polarity identifies a network of cellular programs that control human cancers. Science, 381(6657):515–524, 2023.

Michael P Plebanek, Debayan Bhaumik, Paul J Bryce, and C Shad Thaxton. Scavenger receptor type b1 and lipoprotein nanoparticle inhibit myeloid-derived suppressor cells. Molecular cancer therapeutics, 17(3):686–697, 2018.

Amy R Dwyer, Eloise L Greenland, and Fiona J Pixley. Promotion of tumor invasion by tumor-associated macrophages: the role of csf-1-activated phosphatidylinositol 3 kinase and src family kinase motility signaling. Cancers, 9(6):68, 2017.

Xisong Liang, Zeyu Wang, Ziyu Dai, Hao Zhang, Jian Zhang, Peng Luo, Zaoqu Liu, Zhixiong Liu, Kui Yang, Quan Cheng, et al. Glioblastoma glycolytic signature predicts unfavorable prognosis, immunological heterogeneity, and eno1 promotes microglia m2 polarization and cancer cell malignancy. Cancer Gene Therapy, 30(3):481–496, 2023.

Tanapat Palaga, Wipawee Wongchana, and Patipark Kueanjinda. Notch signaling in macrophages in the context of cancer immunity. Frontiers in immunology, 9:652, 2018.

Shanze Chen, Abdullah FUH Saeed, Quan Liu, Qiong Jiang, Haizhao Xu, Gary Guishan Xiao, Lang Rao, and Yanhong Duo. Macrophages in immunoregulation and therapeutics. Signal Transduction and Targeted Therapy, 8(1):207, 2023.

Haixia Xu, Jimmy Zhu, Sinead Smith, Julia Foldi, Baohong Zhao, Allen Y Chung, Hasina Outtz, Jan Kitajewski, Chao Shi, Silvio Weber, et al. Notch–rbp-j signaling regulates the transcription factor irf8 to promote inflammatory macrophage polarization. Nature immunology, 13(7):642–650, 2012.

Julia Foldi, Yingli Shang, Baohong Zhao, Lionel B Ivashkiv, and Xiaoyu Hu. Rbp-j is required for m2 macrophage polarization in response to chitin and mediates expression of a subset of m2 genes. Protein & cell, 7(3):201–209, 2016.

Tobias Friedrich, Francesca Ferrante, Léo Pioger, Andrea Nist, Thorsten Stiewe, Jean-Christophe Andrau, Marek Bartkuhn, Benedetto Daniele Giaimo, and Tilman Borggrefe. Notch-dependent and-independent functions of transcription factor rbpj. Nucleic Acids Research, 50(14):7925–7937, 2022.

Ruth A Franklin, Will Liao, Abira Sarkar, Myoungjoo V Kim, Michael R Bivona, Kang Liu, Eric G Pamer, and Ming O Li. The cellular and molecular origin of tumor-associated macrophages. Science, 344(6186):921–925, 2014.

Min Liu, Zan Tong, Chuanlin Ding, Fengling Luo, Shouzhen Wu, Caijun Wu, Sabrin Albeituni, Liqing He, Xiaoling Hu, David Tieri, et al. Transcription factor c-maf is a checkpoint that programs macrophages in lung cancer. The Journal of clinical investigation, 130(4):2081–2096, 2020.

Ilaria Marigo, Rosalinda Trovato, Francesca Hofer, Vincenzo Ingangi, Giacomo Desantis, Kevin Leone, Francesco De Sanctis, Stefano Ugel, Stefania Cané, Anna Simonelli, et al. Disabled homolog 2 controls prometastatic activity of tumor-associated macrophages. Cancer discovery, 10(11):1758–1773, 2020.

Shujing Wang, Jingrui Wang, Zhiqiang Chen, Jiamin Luo, Wei Guo, Lingling Sun, and Lizhu Lin. Targeting m2-like tumor-associated macrophages is a potential therapeutic approach to overcome antitumor drug resistance. NPJ Precision Oncology, 8(1):31, 2024.

George C Linderman, Jun Zhao, Manolis Roulis, Piotr Bielecki, Richard A Flavell, Boaz Nadler, and Yuval Kluger. Zero-preserving imputation of single-cell rna-seq data. Nature communications, 13(1):192, 2022.

Etienne Becht, Leland McInnes, John Healy, Charles-Antoine Dutertre, Immanuel WH Kwok, Lai Guan Ng, Florent Ginhoux, and Evan W Newell. Dimensionality reduction for visualizing single-cell data using umap. Nature biotechnology, 37(1):38–44, 2019.

Xinlei Mi, Baiming Zou, Fei Zou, and Jianhua Hu. Permutation-based identification of important biomarkers for complex diseases via machine learning models. Nature communications, 12(1):3008, 2021.

Luyi Tian, Xueyi Dong, Saskia Freytag, Kim-Anh Lê Cao, Shian Su, Abolfazl JalalAbadi, Daniela Amann-Zalcenstein, Tom S Weber, Azadeh Seidi, Jafar S Jabbari, et al. Benchmarking single cell rna-sequencing analysis pipelines using mixture control experiments. Nature methods, 16(6):479–487, 2019.

Amnon Shashua and Tamir Hazan. Non-negative tensor factorization with applications to statistics and computer vision. In Proceedings of the 22nd international conference on Machine learning, pages 792–799, 2005.

Robin Browaeys, Wouter Saelens, and Yvan Saeys. Nichenet: modeling intercellular communication by linking ligands to target genes. Nature methods, 17(2):159–162, 2020.

Christian D Sigg and Joachim M Buhmann. Expectation-maximization for sparse and non-negative pca. In Proceedings of the 25th international conference on Machine learning, pages 960–967, 2008.

Lawrence Page. The pagerank citation ranking: Bringing order to the web. Technical report, Technical Report, 1999.

Andrew Butler, Paul Hoffman, Peter Smibert, Efthymia Papalexi, and Rahul Satija. Integrating single-cell transcriptomic data across different conditions, technologies, and species. Nature biotechnology, 36(5):411–420, 2018.

Christian H Holland, Jovan Tanevski, Javier Perales-Patón, Jan Gleixner, Manu P Kumar, Elisabetta Mereu, Brian A Joughin, Oliver Stegle, Douglas A Lauffenburger, Holger Heyn, et al. Robustness and applicability of transcription factor and pathway analysis tools on single-cell rna-seq data. Genome biology, 21:1–19, 2020.

William M Rand. Objective criteria for the evaluation of clustering methods. Journal of the American Statistical association, 66(336):846–850, 1971.

Peter J Rousseeuw. Silhouettes: a graphical aid to the interpretation and validation of cluster analysis. Journal of computational and applied mathematics, 20:53–65, 1987.

Guangchuang Yu, Li-Gen Wang, Yanyan Han, and Qing-Yu He. clusterprofiler: an r package for comparing biological themes among gene clusters. Omics: a journal of integrative biology, 16(5):284–287, 2012.

Yufei Xiao, Tzu-Hung Hsiao, Uthra Suresh, Hung-I Harry Chen, Xiaowu Wu, Steven E Wolf, and Yidong Chen. A novel significance score for gene selection and ranking. Bioinformatics, 30(6):801–807, 2014.

Laleh Haghverdi, Florian Buettner, and Fabian J Theis. Diffusion maps for high-dimensional single-cell analysis of differentiation data. Bioinformatics, 31(18):2989–2998, 2015.

Greg Finak, Andrew McDavid, Masanao Yajima, Jingyuan Deng, Vivian Gersuk, Alex K Shalek, Chloe K Slichter, Hannah W Miller, M Juliana McElrath, Martin Prlic, et al. Mast: a flexible statistical framework for assessing transcriptional changes and characterizing heterogeneity in single-cell rna sequencing data. Genome biology, 16:1–13, 2015.

Dénes Türei, Alberto Valdeolivas, Lejla Gul, Nicolás Palacio-Escat, Michal Klein, Olga Ivanova, Márton Ölbei, Attila Gábor, Fabian Theis, Dezso Módos, et al. Integrated intra-and intercellular signaling knowledge for multicellular omics analysis. Molecular systems biology, 17(3):e9923, 2021.

Patrick E Meyer, Kevin Kontos, Frederic Lafitte, and Gianluca Bontempi. Information-theoretic inference of large transcriptional regulatory networks. EURASIP journal on bioinformatics and systems biology, 2007:1–9, 2007.

Ian T Jolliffe. A note on the use of principal components in regression. Journal of the Royal Statistical Society Series C: Applied Statistics, 31(3):300–303, 1982.

Yidi Deng, Jarny Choi, and Kim-Anh Lê Cao. Sincast: a computational framework to predict cell identities in single-cell transcriptomes using bulk atlases as references. Briefings in Bioinformatics, 23(3):bbac088, 2022.

Paul-Christian Bürkner. brms: An R package for Bayesian multilevel models using Stan. Journal of Statistical Software, 80(1):1–28, 2017. doi: 10.18637/jss.v080.i01.

Charles E Aull. The first symmetric derivative. The American Mathematical Monthly, 74(6):708–711, 1967.

William G Jacoby. Loess:: a nonparametric, graphical tool for depicting relationships between variables. Electoral studies, 19(4):577–613, 2000.

Alexander J Smola and Risi Kondor. Kernels and regularization on graphs. In Learning Theory and Kernel Machines: 16th Annual Conference on Learning Theory and 7th Kernel Workshop, COLT/Kernel 2003, Washington, DC, USA, August 24-27, 2003. Proceedings, pages 144–158. Springer, 2003.

Sergio Picart-Armada, Wesley K Thompson, Alfonso Buil, and Alexandre Perera-Lluna. The effect of statistical normalization on network propagation scores. Bioinformatics, 37(6):845–852, 2021.

Austin R Benson, Jason D Lee, Bartek Rajwa, and David F Gleich. Scalable methods for nonnegative matrix factorizations of near-separable tall-and-skinny matrices. Advances in neural information processing systems, 27, 2014.

Haipeng Shen and Jianhua Z Huang. Sparse principal component analysis via regularized low rank matrix approximation. Journal of multivariate analysis, 99(6):1015–1034, 2008.

Lester Mackey. Deflation methods for sparse pca. Advances in neural information processing systems, 21, 2008.

Gabor Csardi and Tamas Nepusz. The igraph software. Complex syst, 1695:1–9, 2006.

Florian Rohart, Benoît Gautier, Amrit Singh, and Kim-Anh Lê Cao. mixomics: An r package for ‘omics feature selection and multiple data integration. PLoS computational biology, 13(11):e1005752, 2017.

Svante Wold, Michael Sjöström, and Lennart Eriksson. Pls-regression: a basic tool of chemometrics. Chemometrics and intelligent laboratory systems, 58(2):109–130, 2001.

Yael Baran, Akhiad Bercovich, Arnau Sebe-Pedros, Yaniv Lubling, Amir Giladi, Elad Chomsky, Zohar Meir, Michael Hoichman, Aviezer Lifshitz, and Amos Tanay. Metacell: analysis of single-cell rna-seq data using k-nn graph partitions. Genome biology, 20:1–19, 2019.

